# Genetic nurture effects on education: a systematic review and meta-analysis

**DOI:** 10.1101/2021.01.15.426782

**Authors:** Biyao Wang, Jessie R. Baldwin, Tabea Schoeler, Rosa Cheesman, Wikus Barkhuizen, Frank Dudbridge, David Bann, Tim T. Morris, Jean-Baptiste Pingault

## Abstract

Child educational development is associated with major psychological, social, economic and health milestones throughout the life course. Understanding the early origins of educational inequalities and their reproduction across generations is therefore crucial. Recent genomic studies provide novel insights in this regard, uncovering “genetic nurture” effects, whereby parental genotypes influence offspring’s educational development via environmental pathways rather than genetic transmission. These findings have yet to be systematically appraised. We conducted the first systematic review and meta-analysis to quantify genetic nurture effects on educational outcomes and investigate key moderators. Twelve studies comprising 38,654 distinct parent(s)-offspring pairs or trios from eight cohorts were included, from which we derived 22 estimates of genetic nurture effects. Multilevel random effects models showed that the effect of genetic nurture on offspring’s educational outcomes (*β*_genetic nurture_ = 0.08, 95% CI [0.07, 0.09]) was about half the size of direct genetic effects (*β*_direct genetic_ = 0.17, 95% CI [0.13, 0.20]). Maternal and paternal genetic nurture effects were similar in magnitude, suggesting comparable roles of mothers and fathers in determining their children’s educational outcomes. Genetic nurture effects were largely explained by parental educational level and family socioeconomic status, suggesting that genetically influenced environments play an important role in shaping child educational outcomes. Even after accounting for genetic transmission, we provide evidence that environmentally mediated parental genetic influences contribute to the intergenerational transmission of educational outcomes. Further exploring these downstream environmental pathways may inform educational policies aiming to break the intergenerational cycle of educational underachievement and foster social mobility.

**Public Significance Statement:** This meta-analysis demonstrates that parents’ genetics influence their children’s educational outcomes through the rearing environments that parents provide. This “genetic nurture” effect is largely explained by family socioeconomic status and parental education level, is similar for mothers and fathers (suggesting that both parents equally shape their children’s educational outcomes) and is about half the size of direct genetic effects on children’s educational outcomes. Interventions targeting such environmental pathways could help to break the intergenerational cycle of educational underachievement and foster social mobility.

## Genetic nurture effects on education: a systematic review and meta-analysis

Educational attainment is defined as the highest education level a person attains. A related construct is educational achievement, which refers to one’s school performance. These two constructs are prospectively associated with major psychological, social, economic and health milestones throughout the life-course (Conti, Heckman, & Urzua, 2010; Crespo, López-Noval, & Mira, 2014; Foundation, Kaplan, House, Schoeni, & Pollack, 2008). Parents’ educational level is an important early predictor of their offspring’s own educational attainment and achievement (e.g., Dubow, Boxer, & Huesmann, 2009). It is crucial to understand the processes underlying this transmission of educational attainment and achievement, which leads to continuing cycles of disadvantage across generations and hinders social mobility.

## Origins of Parent-offspring Resemblance on Education

Positive associations between parents’ education and their offspring’s education are found in nearly every society (Björklund & Salvanes, 2011). For example, correlations between parents’ and offspring’s educational outcomes were consistent across twelve Western countries with estimates ranging from *r* = 0.30 (Denmark) to 0.46 (U.S.) (Hertz et al., 2008). The parent-offspring resemblance in educational outcomes can be attributed to nature (genetic variants that offspring inherit from their parents) and nurture (the environment that parents provide for their offspring)(Koellinger & Harden, 2018). These nature and nurture effects are complex and intertwined. For example, the environment created by parents can be partly shaped by genetic influences; parents with a higher genetic propensity for learning may have a greater interest in activities such as reading that, in turn, nurture learning in their offspring. The term “genetic nurture” is used to describe the phenomenon by which parental nature (i.e., parental genotype) influences offspring outcomes by shaping the way that parents nurture (Kong et al., 2018). Genetic nurture effects can therefore be considered indirect effects from parental genotype to offspring’s outcomes that are mediated through the family environment, while nature effects can be considered direct effects through genetic inheritance. Importantly, the direct genetic transmission can generate correlations between parental and child educational outcomes in the absence of any effect of parental nurture in shaping child outcomes (a phenomenon akin to passive-gene environment correlation). Conversely, genetic nurture effects cannot exist in the absence of a nurturing effect. This is because parental genotypes that are not transmitted to children can only affect child outcomes via environmental pathways. As such, detecting significant genetic nurture effects provides strong evidence that environmental pathways matter when it comes to shaping children’s educational outcomes, even after accounting for genetic transmission.

## Molecular Genetic Evidence of Environmentally Mediated Effects on Education

Findings from genome-wide association studies (GWAS) in the last two decades have greatly advanced our understanding of complex traits, including educational outcomes (Visscher et al., 2017). The most recent GWAS for educational attainment (hereafter referred to as “EA GWAS”) identified 1271 lead variants associated with year of schooling completed (Lee et al., 2018). Based on the EA GWAS, a polygenic score (PGS) can be derived to provide a single value reflecting an individual’s genetic propensity to educational attainment (referred to as “EA PGS”; it is a sum of an individual’s effect alleles weighted by effect sizes obtained from the EA GWAS)(Dudbridge, 2013; Ronald, 2020). The top performing EA PGS explained 11-13% of the variance in educational attainment in two replication samples (Lee et al., 2018).

Early studies have shown that the EA PGS relates to educational outcomes in parents and offspring, highlighting the potential role that genetics may play in the intergenerational transmission of educational outcomes (Ayorech, Krapohl, Plomin, & von Stumm, 2017; Belsky et al., 2016; Domingue, Belsky, Conley, Harris, & Boardman, 2015; Krapohl & Plomin, 2016; Selzam et al., 2017). More advanced designs were recently devised to further disentangle direct versus indirect pathways underlying the intergenerational transmission of educational outcomes. Two studies (Bates et al., 2018; Kong et al., 2018) assessed the magnitude of genetic nurture effects by constructing PGS from parental alleles that are not transmitted to the offspring. This approach is termed the “*virtual parent design*” (for more description see section 1.1 in the supplement) as it creates a ‘virtual parent’ that is not genetically related to the offspring. Hence, the association of the non-transmitted parental PGS with offspring outcomes occurs not though genetic transmission but environmental pathways. The effect of non-transmitted PGS on offspring outcomes therefore reflect genetic nurture effects by design because it is free from genetic confounding between parents and offspring due to shared genotype. Moreover, as the transmitted genotypes from both parents (i.e., the inherited offspring genotype) can have both direct and genetic nurture effects, the association between child PGS and their own outcomes can be overestimated when genetic nurture is not considered (Kong et al., 2018). Direct genetic effects represent genetic influences that originate in the child genetic variants and are corrected for the inflation arising from genetic nurture. Previous findings suggested that genetic nurture has substantial impacts on offspring’s educational attainment, while it is less important for other traits such as body mass index and height (Kong et al., 2018). This design has now been implemented in different contexts, such as using cohorts from different countries, constructing PGS based on more powerful/accurate GWAS, etc (e.g., Bates et al., 2019; de Zeeuw et al., 2020).

In addition to relying on non-transmitted and transmitted polygenic scores, genetic nurture and direct genetic effects can be obtained by estimating the effect of parental PGS on the offspring’s outcome, while statistically controlling for the offspring’s PGS (for more description see section 1.2 in the supplement). This statistical control approach has been applied in several studies (e.g., Armstrong-Carter et al., 2020; T. Morris, Davies, Hemani, & Smith, 2020; Willoughby, McGue, Iacono, Rustichini, & Lee, 2019). Estimating genetic nurture effects using statistical control requires genetic data on offspring and both parents to obtain unbiased estimates, but can provide an approximation of the genetic nurture effects when genetic data is available for one parent only.

## Sources of Heterogeneity in Genetic Nurture Effects

The magnitude of genetic nurture effects on children’s educational outcomes may vary according to four key factors, namely (1) parent of origin, (2) analytic method (3) outcome type (4) and predictive accuracy of the GWAS used to derive the PGS. First, it is unclear whether the impact of genetic nurture differs depending on whether it originates in the mother or the father. While, historically, mothers might have been expected to have a greater “genetic nurturing” impact on their children (assuming a greater involvement in nurturing their offspring), emerging studies from modern contexts have emphasized the important role of fathers in children’s development (Barger, Kim, Kuncel, & Pomerantz, 2019; Kim & Hill, 2015). However, previous studies have rarely directly compared effects of maternal versus paternal genetic nurture on children’s educational outcomes. Second, it is unclear whether the magnitude of genetic nurture effects in empirical studies differ depending on the analytic method used (i.e., virtual parent or statistical control). Moreover, due to lack of complete trio data (i.e. child and both parents), genetic nurture effects have often been estimated among parent-offspring pairs. It is unclear to what extent the missing parental genotype bias estimates. Third, it is unclear whether the effect of genetic nurture differs depending on the type of educational outcome assessed (e.g., educational attainment or achievement). Previous genetically informed studies on educational outcomes have focused on attainment or achievement interchangeably (e.g., Allegrini et al., 2019; Krapohl & Plomin, 2016; Sorensen et al.; von Stumm et al., 2020). However, as both efforts relied on the GWAS of educational attainment, which is likely to more strongly correlate with educational attainment than achievement at the phenotypic level, studies examining educational attainment may capture genetic nurture effects more accurately. Fourth, to what extent the estimated genetic nurture effects differ depending on the accuracy of GWAS summary statistics used to derive the PGS remains untested. Studies have used different GWAS data depending on the most powerful data at that time of publication. For example, the number of detected lead genetic variants increased from three in the first (Rietveld et al., 2013) to 1,271 in the most recent (Lee et al., 2018) GWAS of EA, thereby increasing the ability to explain variance in children’s educational outcomes from 2% to 11% (additional details in section 2 of the supplement).

## Summary and Aim

Here, we conduct the first meta-analysis of all PGS studies on educational outcomes in parent(s)-offspring samples published to date to answer the following questions: (1) What is the magnitude of genetic nurture effects on educational outcomes? (2) What is the magnitude of direct genetic effects on educational outcomes when accounting for genetic nurture effects? (3) Which factors moderate genetic nurture and direct genetic effects on educational outcomes?

## Method

### Search Strategy and Study Selection

This systematic review and meta-analysis was performed in line with the Preferred Reporting Items for Systematic Reviews and the Meta-Analyses (PRISMA (Moher, Liberati, Tetzlaff, Altman, & Group, 2009)) statement and Meta-Analyses of Observational Studies in Epidemiology (MOOSE (Stroup et al., 2000)) guidelines (checklists in sTables 1 and 2 in the supplement). The protocol was registered on the Open Science Framework (https://osf.io/q8b25/). The literature search was performed in July 2020 in Ovid (MEDLINE, EMBASE, PsycINFO), Web of Science Core Collection, PubMed for peer-reviewed articles written in English. To estimate genetic nurture effects on educational outcomes, we considered articles estimating parental genetic nurture using EA PGS. Therefore, the publication period was limited to 2013 onwards, when the first EA GWAS (Rietveld et al. 2013) became available. To retrieve relevant publications, the search included terms related to: (1) educational outcomes, (2) polygenic scores, and (3) genetic nurture effects. Detailed literature search strategy and terms are presented in section 3.1 in the supplement. Two authors (B.W. and T.S.) independently screened titles and abstracts of all articles retrieved from the search before reviewing the full text of potentially eligible studies (see criteria below). Disagreements were resolved through discussion with the senior researcher (J.B.P). Eligible studies met the following criteria: (1) They assessed offspring educational attainment (e.g., years of education, highest degree obtained) or educational achievement (e.g., national tests scores or levels, school grades) in the general population. (2) The exposure variable(s) included genomic proxies of education in parents and offspring, measured in the form of PGS derived from the EA GWAS. (3) Studies derived estimates for genetic nurture effects on education based on one of the following designs that rely on genotype data from parents and their biological offspring: (a) virtual parent: testing whether the PGS calculated from parents’ non-transmitted alleles predict offspring educational outcomes; or (b) statistical control: testing whether parents’ PGS predict offspring educational outcomes over and above offspring’s own PGS. For more information on inclusion criteria see section 3.2 in the supplement.

**Table 1.** Studies investigating genetic nurture effects on educational outcomes

| Cohort<sup>a</sup> | Publication | Outcome<sup>b</sup> | Effective<br>N<sup>c</sup> | Design | GWAS<sup>d</sup> | NOS<br>score<sup>e</sup> |
| --- | --- | --- | --- | --- | --- | --- |
| Born in Bradford birth cohort (BiB), United Kindom | Armstrong-Carter et al., 2020 | Key stage 1 school-based exam score | 1267 | Statistical control | EA3 | 6.5 |
| The Brisbane Adolescent Twin Study (BATS), Australia | Bates et al., 2018 | The Queensland Core Skills Test | 2335 | Virtual parent | EA2 | 7.5 |
| The Brisbane Adolescent Twin Study (BATS), Australia | Bates et al., 2019 | The Queensland Core Skills Test | 2335 | Virtual parent | EA3 | 7.5 |
| The Environmental Risk Longitudinal Twin Study (E-Risk), United Kingdom | Belsky et al., 2018 | GCSE academic qualification level | 1574 | Statistical control | EA3 | 7.0 |
| The Framingham Heart Study (FHS), United States | Conley et al., 2015 | Years of schooling | 968 | Statistical control | EA1 | 5.0 |
| The Netherlands Twin Register (NTR), Netherlands | de Zeeuw et al., 2020 | Highest obtained degree; Nationwide educational achievement test | 1931;<br>1120 | Virtual parent | EA3 | 7.0 |
| The Icelandic quantitative trait cohorts (deCODE), Iceland | Kong et al., 2018 | Years of education completed | 21637 | Virtual parent | EA2 | 7.5 |
| The Framingham Heart Study (FHS), United States | Liu et al., 2018 | Years of education completed | 6298 | Statistical control | EA2 | 6.5 |
| The Avon Longitudinal Study of Parents and Children (ALSPAC), United Kingdom | Morris 2020 | Key stage 4 school-based exam score | 1095 | Statistical control | EA3 | 7.0 |
| The Minnesota Twin Family Study (MTFS), United States | Rustichini et al., 2018 | Years of education completed; High school grades | 1690;<br>1583 | Statistical control | EA3 | 6.0 |
| The Environmental Risk Longitudinal Twin Study(E-Risk),<br>United Kingdom | Wertz et al., 2019 | GCSE academic<br>qualification level | 860 | Statistical control | EA3 | 6.0 |
| The Minnesota Center for Twin and Family Research<br>(MCTFR), United States | Willoughy et al., 2019 | High school grades<sup>a</sup> | 2517 | Statistical control | EA3 | 5.5 |

*Note.* <sup>a</sup> Participants in the MCTFR cohort were drawn from several longitudinal studies including the MTFS cohort, thus in the meta-analysis they were considered as the same cohort. <sup>b</sup> Educational outcomes consists of two broad categories, i.e., attainment and achievement. Years of schooling/education completed and highest obtained degree are categorized as educational attainment; the rest are categorized as educational achievement. More details of outcomes, including assessing time, are reported in sTable 3 in the supplement. <sup>c</sup> The largest sample size used to assess genetic nurture effects. <sup>d</sup> GWAS (genome-wide association studies) used to derive the polygenic scores, including EA1 with N = 101,069 (Rietveld et al., 2013), EA2 with N = 293,723 (Okbay et al., 2016), EA3 with N = 1,131,881 (Lee et al., 2018). <sup>e</sup> Quality score ranged from 0 (lowest) to 9 (highest) on methodological quality using an adjusted version of the Newcastle–Ottawa scale, criteria showed in sMethods and detailed scoring showed in sTable 4 in the supplementary material.

**Table 2.** Three-level random effects models of parental and child effects on educational outcomes

| Model | MREM of parental effects |  |  | MREM of child effects |  |  |
| --- | --- | --- | --- | --- | --- | --- |
|  | Mixture<sup>a</sup> | Genetic nurture effects | Unadjusted parental effects | Mixture<sup>b</sup> | Direct genetic effects | Unadjusted child effects |
| k<sub>cohort</sub> | 8 | 8 | 3 | 8 | 8 | 7 |
| k<sub>estimate</sub> | 30 | 22 | 8 | 27 | 16 | 11 |
| $\beta_{\text{pooled}}$ | 0.11 | 0.08 | 0.21 | 0.20 | 0.17 | 0.24 |
| $\beta_{95\% \text{ CI}}$ | 0.08-0.14 | 0.07-0.09 | 0.15-0.26 | 0.16-0.24 | 0.13-0.20 | 0.20-0.28 |
| $\beta_{\text{robust CI}}^c$ | 0.07-0.14 | 0.06-0.10 | 0.08-0.33 | 0.15-0.24 | 0.12-0.21 | 0.19-0.29 |
| $\sigma^2_{\text{Level 2}}$ | $\chi^2 = 45.23, p < .0001$ | $\chi^2 < 0.01, p = .5000$ | $\chi^2 < 0.01, p = .5000$ | $\chi^2 = 60.17, p < .0001$ | $\chi^2 < 0.01, p = .5000$ | $\chi^2 = 1.55, p = .1067$ |
| $\sigma^2_{\text{Level 3}}$ | $\chi^2 = 1.23, p = .1339$ | $\chi^2 = 1.94, p = .0817$ | $\chi^2 = 5.73, p = .0083$ | $\chi^2 = 5.46, p = .0097$ | $\chi^2 = 5.09, p = .0120$ | $\chi^2 = 1.27, p = .1298$ |
| $I^2_{\text{Level 1}}$ | 10.17% | 76.80% | 19.14% | 8.70% | 17.67% | 11.57% |
| $I^2_{\text{Level 2}}$ | 65.83% | <0.01% | <0.01% | 36.17% | <0.01% | 28.51% |
| $I^2_{\text{Level 3}}$ | 24.00% | 23.20% | 80.86% | 55.13% | 82.33% | 59.92% |
| Publ, bias | $Q = 4.01, p = .0453$ | $Q = 6.12, p = .0134$ | $Q = 3.22, p = .0727$ | $Q = 9.56, p = .0020$ | $Q = 0, p = .9976$ | $Q = 0.47, p = 0.4917$ |

Note. <sup>a</sup> Effect sizes of genetic nurture and unadjusted parental effects were jointly modelled. <sup>b</sup> Effect sizes of direct genetic and unadjusted child effects were jointly modelled.
<sup>c</sup> Robust confidence intervals were cluster-robust variance estimations not estimated from multilevel random effects model, for details see section 7.1 in the supplement.
MREM = Multilevel random effects model; $\beta$ = standardized regression coefficients (i.e., the metric of effect sizes); CI = confidence interval; $\chi^2$ Statistics from likelihood-ratio test to test within-cohort variance ( $\sigma^2_{\text{Level 2}}$ ) and between-cohort variance ( $\sigma^2_{\text{Level 3}}$ ) for significance; $I^2$ = % of the total variance accounted for by random sampling variance (Level 1), variation within cohorts (Level 2), variation between cohorts (Level 3); Publi. Bias = publication bias, assessed by using precision (sampling variance) to predict the effect size.

### Quality Assessment, Data Extraction and Effect Size Calculation

The methodological quality of the included studies was assessed by two authors (among B.W., J.B., W.B. and R.C.) using an adapted version of the Newcastle–Ottawa scale (NOS (Stang, 2010)). The NOS was adapted for its use on genetically informed studies and included nine questions tapping into four wider aspects relevant to study quality, including the quality of cohort selection, the assessment of exposure, the level of comparability of the cohort, and the assessment of outcomes. Overall study quality was indexed as a sum score ranging from 0 to 9 (see section 3.3 for detailed scoring criteria and sTable 4 for score of each study in the supplement).

Data extraction for each included study was independently performed by two of the authors (among B.W., J.B., W.B. and R.C.). The following data were extracted: publication characteristics (study name, first author, year), sample characteristics (cohort name, sample size, population source, ethnicity, sex distribution), study design (virtual parent or statistical control), calculation of PGS (GWAS used to derive the PGS, PGS threshold, source/parent of origin of genotype, whether standardized), education-related outcome assessed (educational outcome, outcome type, age at assessment, whether standardized), effect size (estimation type, estimation, 95% CI or standard error of the estimation), and adjusted confounding variables. Where information was missing, original study authors were contacted to request the information.

As a common metric, we extracted (or converted effect sizes to) standardized beta coefficients and corresponding standard errors from all individual studies. These data were then included in our meta-analytical models to derive the pooled estimate of genetic nurture effects. For studies using the virtual parent design, we extracted standardized regression coefficients of non-transmitted PGS. For studies using the statistical control design, we extracted adjusted standardized regression coefficients of parental PGS, while controlling for offspring’s PGS. For studies reporting effect estimates in metrics other than standardized beta or without corresponding standard errors, we transformed the reported statistics using the formulae included in the R package compute.es_0.2-4 (Del Re, 2013). One estimate of genetic nurture derived from an average parental PGS was recalibrated to be comparable with other studies using PGS of individual parent (for justification see section 7.2 in the supplement). Estimates of direct genetic effects were extracted when available or imputable, i.e., the difference between standardized regression coefficients of transmitted PGS and non-transmitted PGS in the virtual parent design or adjusted standardized regression coefficients of offspring’s PGS while statistically controlling for parental PGS. Whenever applicable, we also derived unadjusted parental or child effects, namely regression coefficients of offspring’s educational outcomes on parental or offspring’s PGS. For more information on the effect size transformation and calculation see section 4.1 in the supplement. With each article reviewed and coded by two authors, the two coders had interrater reliabilities of 92.6% on quality assessment and 97.8% on data extraction. Before moving onto analyses, discrepancies were reviewed and arbitrated by the two coders, and disagreements were resolved through discussion with the senior researcher (J.B.P).

### Statistical Analysis

Analyses were conducted in R version 3.6.1 (R Core Team, 2019) using the package metafor version 2.4-0 (Viechtbauer, 2010). Since multiple effect sizes were reported in individual studies and cohorts, we used three-level Multilevel Random-Effects Models (MREMs) to account for dependency among effect sizes within single studies/cohorts (i.e., correlation between effect sizes). These models incorporate three variance components, namely sampling variance at level 1 (variance that is unique for each estimated effect size), within-cohort variance at level 2 (variance across outcomes within a cohort), and between-cohort variance at level 3 (variance across cohorts). For more information on multilevel random-effects models see section 4.2 in the supplement. We assessed the heterogeneity between studies using the *I*^2^ statistics and tested whether heterogeneity of effect sizes at level 2 (within-cohort heterogeneity) and level 3 (between-cohort heterogeneity) was significant by conducting two separate one-sided log-likelihood ratio tests (Assink & Wibbelink, 2016). Publication bias were visualized by checking the asymmetry of funnel plots and more formally tested by using precision (sampling variance) as a moderator in meta-analysis models (Rodgers & Pustejovsky, 2020).

Meta-regression analyses were performed to explore potential sources of heterogeneity in effect sizes. We tested four main categorical moderators: (1) The type of parental genotype used for constructing PGS, which can be maternal or paternal genotype, or the mixture of both. (2) The type of analytic method used to measure the genetic nurture effects, which can be virtual parent, partial or full statistical control. (3) The type of educational outcome assessed, which can be educational attainment or educational achievement. (4) The specific GWAS summary statistics used to derive PGS, which can be EA1 with N = 101,069 (Rietveld et al., 2013), EA2 with N = 293,723 (Okbay et al., 2016), EA3 with N = 1,131,881 (Lee et al., 2018). In addition, we tested the moderating role of study characteristics (i.e., methodological quality, sample size and attrition in cohort), for details see section 4.3 in the supplement. We also tested whether adjusting for observed parental educational level and family socioeconomic status (SES) attenuated genetic nurture (details in section 6 in the supplement).

We undertook a series of sensitivity checks to evaluate the robustness of our results including: computing robust confidence intervals, evaluating the impact of recalibrating effects derived from average parental PGS in one study (Willoughby et al., 2019), assessing the impact of a potentially influential study (Kong et al., 2018), performing jack-knife leave-one-out analyses and assessing the moderating effect of outcome type within study (i.e., when educational attainment and achievement were measured in the same study). For more information on sensitivity analyses, see section 7 in the supplement. In all tests, a 2-tailed *p* < .05 was considered statistically significant.

## Results

### Study Description

Twelve studies met the inclusion criteria (see Figure 1 for study selection procedure and Table 1 for study summary. Further details see sTable 3 in the supplement). The studies comprised 38,654 distinct offspring individuals (computation of total sample size see section 4.3 in the supplement) plus at least one of their parents across eight study cohorts. We derived *k* = 22 estimates of genetic nurture effects on educational outcomes and *k* = 16 estimates of direct genetic effects. Additionally, estimates of unadjusted parental (*k* = 8) and child (*k* = 11) effects were extracted. The majority of genetic nurture estimates were derived from studies using the statistical control approach [68.2% (*k* = 15)] and the rest from virtual parent design [31.8% (*k* = 7)]. Slightly more studies focused on educational achievement [54.5% (*k* = 12)] versus educational attainment [45.5% (*k* = 10)].

**Table 3.**
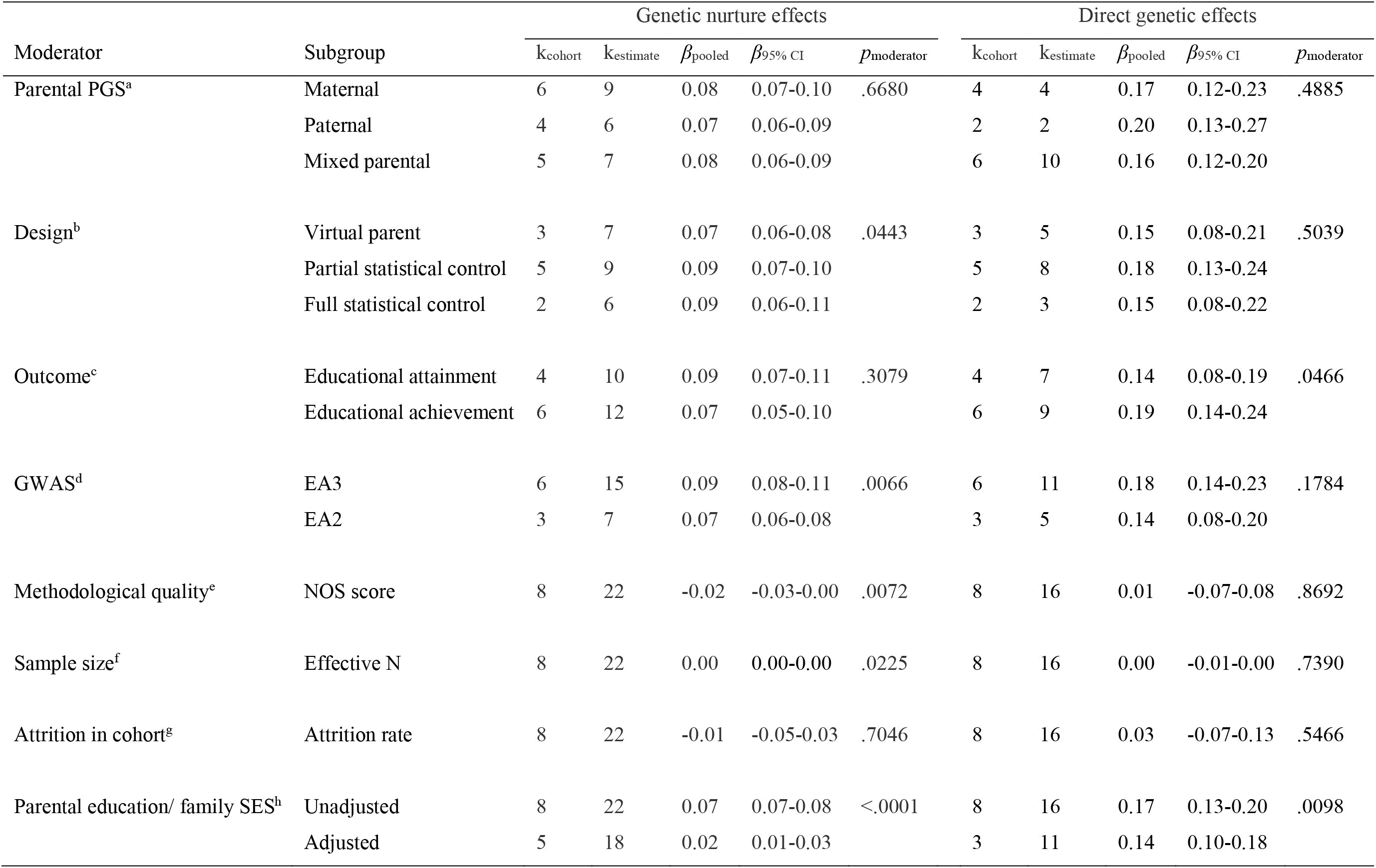
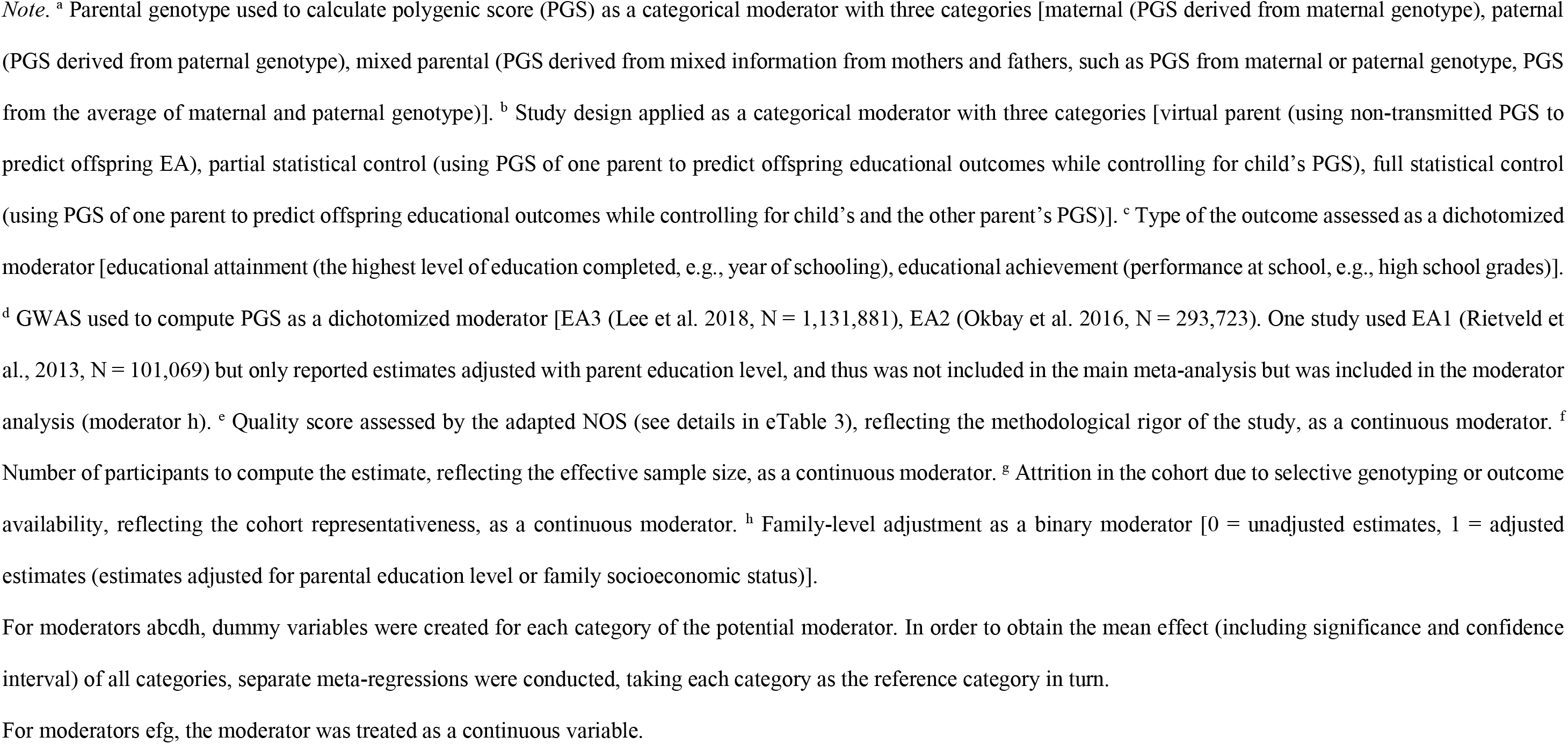
moderator analysis: sources of heterogeneity in MREM of genetic nurture effects and direct genetic effects

| Moderator | Subgroup | Genetic nurture effects |  |  |  |  | Direct genetic effects |  |  |  |  |
| --- | --- | --- | --- | --- | --- | --- | --- | --- | --- | --- | --- |
| | | k<sub>cohort</sub> | k<sub>estimate</sub> | $\beta_{\text{pooled}}$ | $\beta_{95\% \text{ CI}}$ | $p_{\text{moderator}}$ | k<sub>cohort</sub> | k<sub>estimate</sub> | $\beta_{\text{pooled}}$ | $\beta_{95\% \text{ CI}}$ | $p_{\text{moderator}}$ |
| Parental PGS<sup>a</sup> | Maternal | 6 | 9 | 0.08 | 0.07-0.10 | .6680 | 4 | 4 | 0.17 | 0.12-0.23 | .4885 |
|  | Paternal | 4 | 6 | 0.07 | 0.06-0.09 |  | 2 | 2 | 0.20 | 0.13-0.27 |  |
|  | Mixed parental | 5 | 7 | 0.08 | 0.06-0.09 |  | 6 | 10 | 0.16 | 0.12-0.20 |  |
| Design<sup>b</sup> | Virtual parent | 3 | 7 | 0.07 | 0.06-0.08 | .0443 | 3 | 5 | 0.15 | 0.08-0.21 | .5039 |
|  | Partial statistical control | 5 | 9 | 0.09 | 0.07-0.10 |  | 5 | 8 | 0.18 | 0.13-0.24 |  |
|  | Full statistical control | 2 | 6 | 0.09 | 0.06-0.11 |  | 2 | 3 | 0.15 | 0.08-0.22 |  |
| Outcome<sup>c</sup> | Educational attainment | 4 | 10 | 0.09 | 0.07-0.11 | .3079 | 4 | 7 | 0.14 | 0.08-0.19 | .0466 |
|  | Educational achievement | 6 | 12 | 0.07 | 0.05-0.10 |  | 6 | 9 | 0.19 | 0.14-0.24 |  |
| GWAS<sup>d</sup> | EA3 | 6 | 15 | 0.09 | 0.08-0.11 | .0066 | 6 | 11 | 0.18 | 0.14-0.23 | .1784 |
|  | EA2 | 3 | 7 | 0.07 | 0.06-0.08 |  | 3 | 5 | 0.14 | 0.08-0.20 |  |
| Methodological quality<sup>e</sup> | NOS score | 8 | 22 | -0.02 | -0.03-0.00 | .0072 | 8 | 16 | 0.01 | -0.07-0.08 | .8692 |
| Sample size<sup>f</sup> | Effective N | 8 | 22 | 0.00 | 0.00-0.00 | .0225 | 8 | 16 | 0.00 | -0.01-0.00 | .7390 |
| Attrition in cohort<sup>g</sup> | Attrition rate | 8 | 22 | -0.01 | -0.05-0.03 | .7046 | 8 | 16 | 0.03 | -0.07-0.13 | .5466 |
| Parental education/ family SES<sup>h</sup> | Unadjusted | 8 | 22 | 0.07 | 0.07-0.08 | <.0001 | 8 | 16 | 0.17 | 0.13-0.20 | .0098 |
|  | Adjusted | 5 | 18 | 0.02 | 0.01-0.03 |  | 3 | 11 | 0.14 | 0.10-0.18 |  |

**Figure 1.**
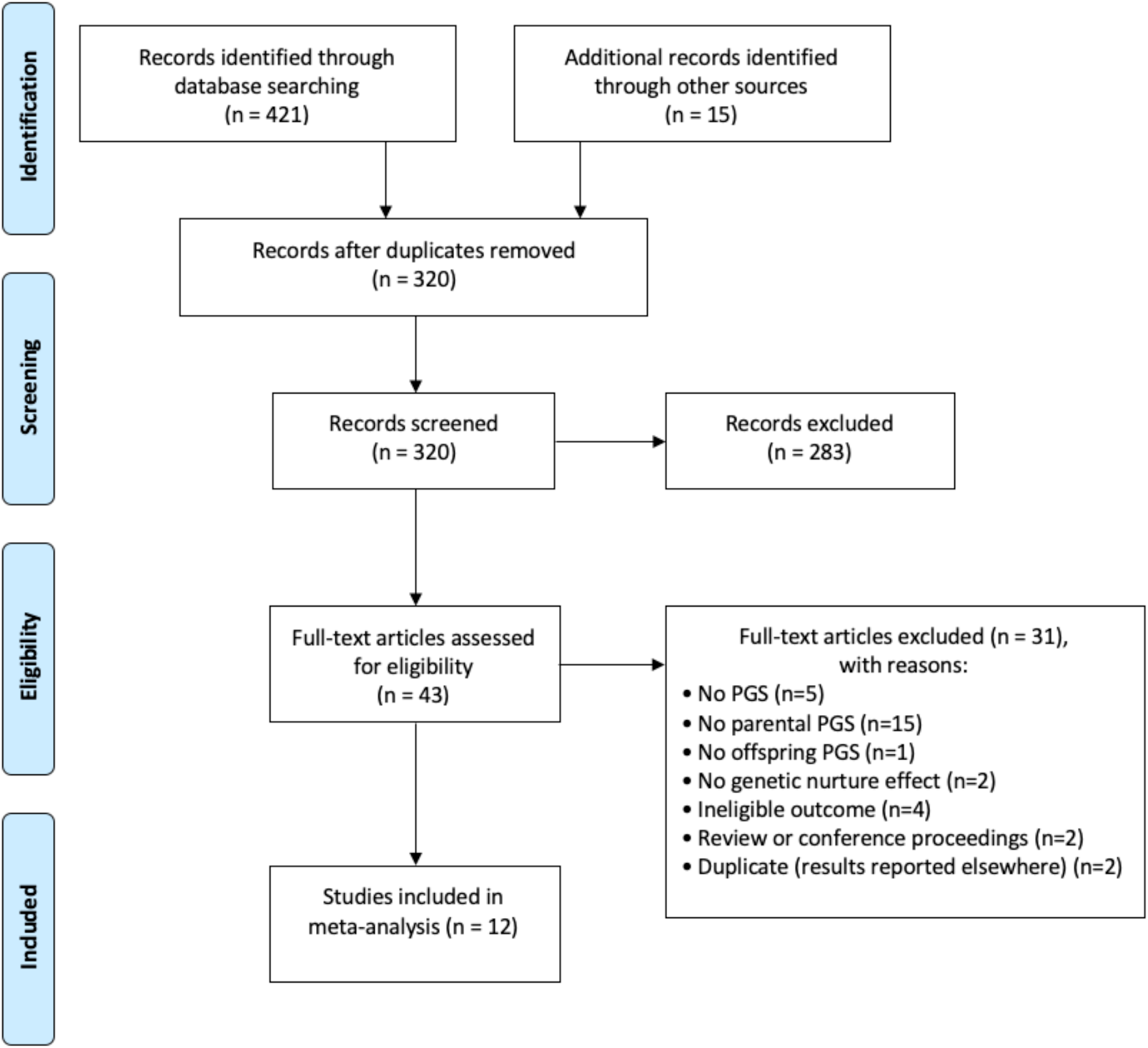
Flowchart

### Genetic nurture effects on offspring’s educational outcomes

As shown in Table 2 and Figure 2, the magnitude of genetic nurture effects was small but precisely estimated (*β*_genetic nurture_ = 0.08, 95% CI [0.07, 0.09], robust CI [0.06, 0.10]). Among estimates of genetic nurture effects, the variance was largely attributed to sampling differences (*I*^2^_Level 1_ = 76.80%). Within-cohort heterogeneity was close to null (*I*^2^_Level 2_ = >0.01%, *χ*^2^_Level 2_ < .01, *p* = .50) and between-cohort heterogeneity was minimal (*I*^2^_Level 3_ = 23.20%, *χ*^2^_Level 3_ = 1.94, *p* = .0817), suggesting largely homogeneous genetic nurture effects across contexts. The funnel plot was visually symmetric (see sFigure 1 in the supplement) but a formal test with precision as the moderator (*Q* = 6.12, *p* = .0134) found some evidence of publication bias in genetic nurture effects. This bias was no longer significant in the sensitivity analysis when excluding the potentially influential study (Kong et al., 2018)(*Q* = 0.88, *p* = .3486, see sTable 5 in the supplement). Results from jack-knife analyses suggest no substantial role of unduly large effect arising from any individual study (see sFigure 2 in the supplement). The supplement includes findings regarding: meta-analytical estimates of unadjusted parental effects and the joint meta-analysis of genetic nurture and unadjusted parental effects (section 5.1), and the impact of recalibrating the average parental PGS (section 7.2).

**Figure 2.**
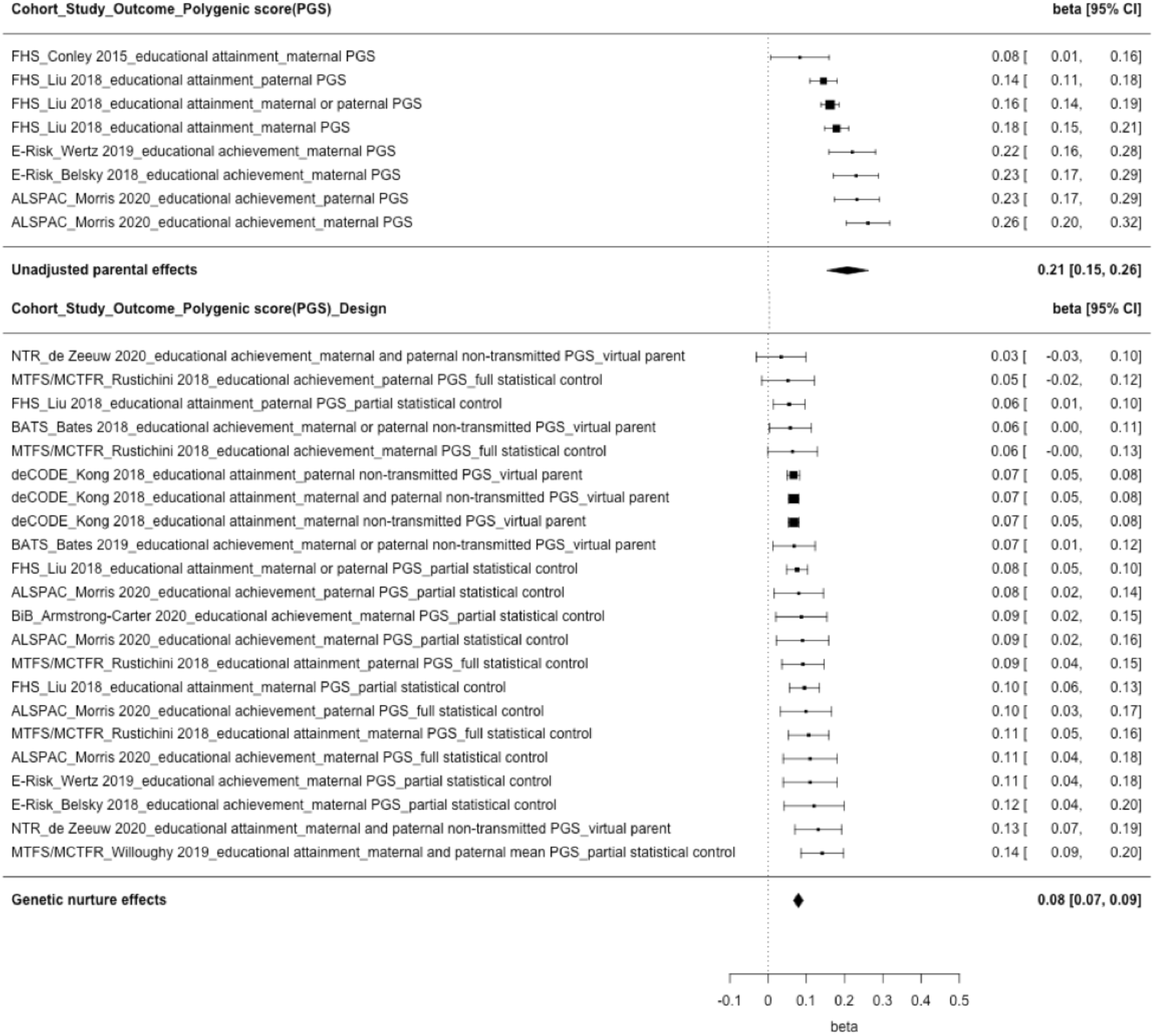
Forest plot of multilevel random effects model for unadjusted parental effects and genetic nurture effects on educational outcomes Note. Effect sizes were standardized beta coefficients, which represent how many standard deviations of change in educational outcome occur per standard deviation of change in EA PGS.

### Direct genetic effects on offspring’s educational outcomes

As shown in Table 2 and Figure 3, the magnitude of direct genetic effects was larger than the genetic nurture effects (*β*_direct genetic_ = 0.17, 95% CI [0.13, 0.20], robust CI [0.12, 0.21]).

**Figure 3.**
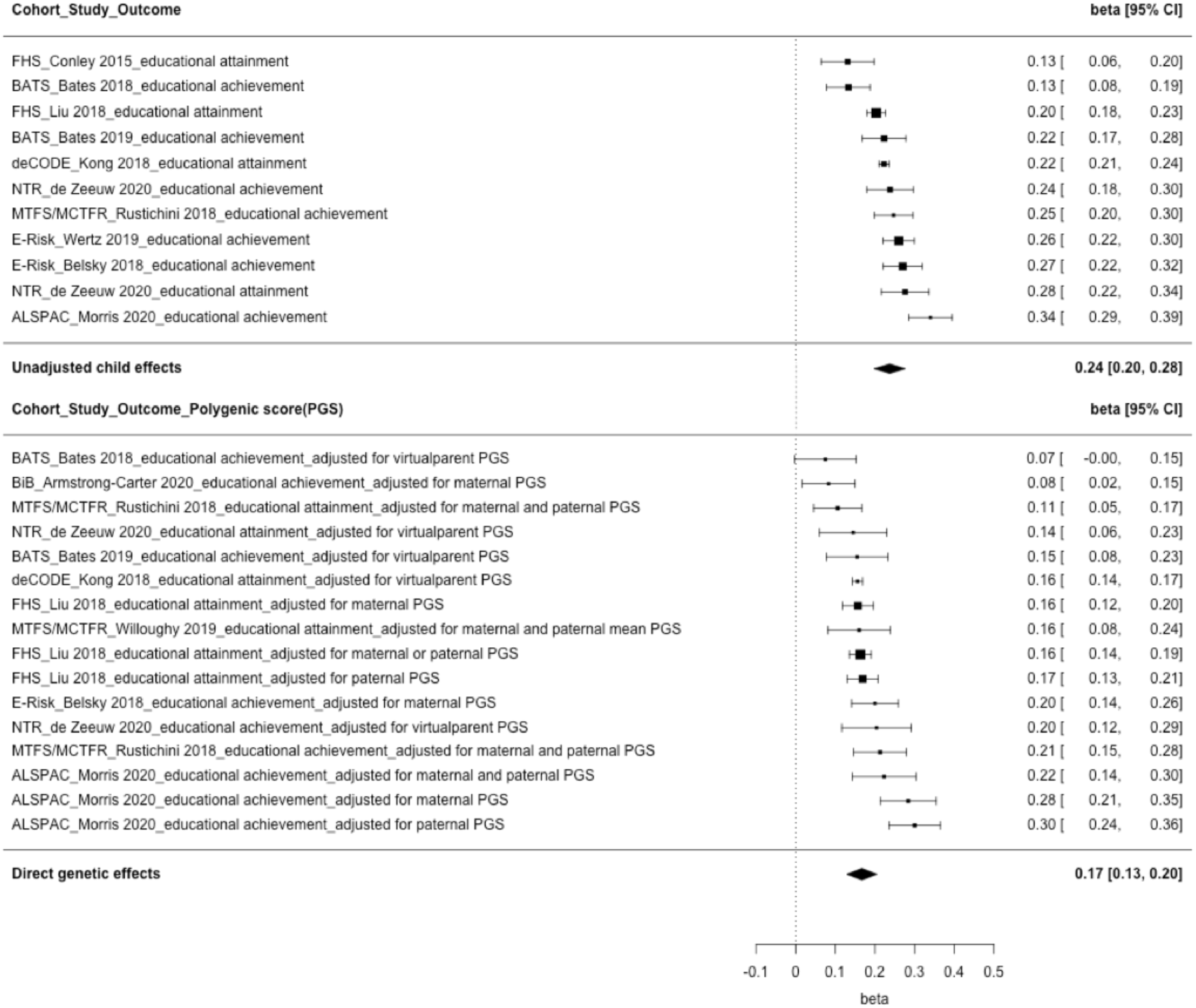
Forest plot of multilevel random effects model for unadjusted child effects and direct genetic effects on educational outcomes Note. Effect sizes were standardized beta coefficients, which represent how many standard deviations of change in educational outcome occur per standard deviation of change in EA PGS.

Among estimates of direct genetic effects, 17.67% (i.e., *I*^2^_Level 1_) of the variance among effect sizes was explained by random sampling. While within-cohort heterogeneity was negligible (*I*^2^_Level 2_ = <0.01%, *χ*^2^_Level 2_ < .01, *p* = .50), the variance was largely attributable to between-cohort heterogeneity (*I*^2^_Level 3_ = 82.33%, *χ*^2^_Level 3_ = 5.09, *p* = .0120). The funnel plot (see sFigure 3 in the supplement) and formal test with precision as a moderator (Table 2) suggested no publication bias in estimates of direct genetic effects. The jack-knife analyses suggested that no single study unduly influenced meta-analysis estimates (see sFigure 4 in the supplement). For findings on unadjusted child effects and the joint meta-analysis of direct genetic and unadjusted child effects, see section 5.2 in the supplement.

### Sources of Heterogeneity in genetic nurture and direct genetic effects on educational outcomes

We tested potential sources of heterogeneity across estimates with meta-regression analysis. Results for genetic nurture and direct genetic effects are displayed in Table 3 (see sTable 6 in the supplement for findings on unadjusted parental and child effects).

#### Parent of Origin

First, we tested whether the parent of origin moderated effect sizes of genetic nurture and direct genetic effects. The MREM suggested similar effects of genetic nurture on educational outcomes when polygenic scores were derived from mothers or fathers (*β*_mother_ = 0.08, 95% CI [0.07, 0.10] for maternal PGS only, *β*_father_ = 0.07, 95% CI [0.06, 0.09] for paternal PGS only, *β*_parents_ = 0.08, 95% CI [0.06, 0.10] for mixed parental PGS, e.g., maternal or paternal PGS, mean of maternal and paternal PGS) as well as direct genetic effects *β*_mother_ = 0.17, 95% CI [0.12, 0.23], *β*_father_ = 0.20, 95% CI [0.13, 0.27], *β*_parents_ = 0.16, 95% CI [0.12, 0.20]). There was no evidence for moderating effects (*p*_genetic nurture_ = .6680 and *p*_direct genetic_ = .4885). These findings were robust to the removal of the potentially influential study (Kong et al., 2018)(see sTable 7 in the supplement).

#### Study Design

Second, we examined whether using different designs moderated effect sizes by comparing estimates relying on virtual parent (using parental non-transmitted PGS to predict children’s education), partial statistical control (using PGS of one parent to predict children’s education while controlling for the child’s PGS) and full statistical control (using PGS of one parent to predict children’s education while controlling for the child’s and the other parent’s PGS). Here, genetic nurture effects detected by the virtual parent design (*β*_virtual parent_ = 0.07, 95% CI [0.06, 0.08]) were lower than those obtained from the statistical control approach (*β*_partial control_ = 0.09, 95% CI [0.07, 0.10], *β*_full control_ = 0.09, 95% CI [0.06, 0.11], *p* = .0443). In contrast, different designs captured similar effect sizes for direct genetic effects (*β*virtual parent = 0.15, 95% CI [0.08, 0.21], *β*_partial control_ = 0.18, 95% CI [0.13, 0.24], *β*_full control_ = 0.15, 95% CI [0.08, 0.22], *p* = .5039).

#### Type of Educational Outcome

Third, we considered whether the type of educational outcome moderated effect sizes by comparing studies assessing educational attainment and educational achievement. Similar effect sizes of genetic nurture effects were found for educational attainment and achievement (*β*_attainment_ = 0.09, 95% CI [0.07, 0.11], *β*_achievement_ = 0.07, 95% CI [0.05, 0.10], *p* = .3079). Direct genetic effects were larger for educational achievement (*β*achievement = 0.19, 95% CI [0.14, 0.24]) than for educational attainment (*β*_attainment_ = 0.14, 95% CI [0.08, 0.19]) and there was evidence of a moderating effect (*p* = .0466). The robustness of this finding was confirmed by restricting the analysis to studies reporting effects for both attainment and achievement (see sTable 8 in the supplement).

#### Predictive accuracy of the GWAS used to derive the PGS

Fourth, we tested whether the accuracy of GWAS used to derive the PGS moderated effect sizes by comparing studies constructing PGS based on different EA GWAS. Estimates of genetic nurture effects based on more accurate GWAS were significantly larger (*β*_EA3_= 0.09, 95% CI [0.08, 0.11], *β*_EA2_= 0.07, 95% CI [0.06, 0.08], *p*_genetic nurture_ = .0066). There was no significant difference for direct genetic effects due to the larger uncertainty in estimates (*β*_EA3_ = 0.18, 95% CI [0.14, 0.23], *β*_EA2_ = 0.14, 95% CI [0.08, 0.20], *p*_direct genetic_ = .1784).

We also considered whether study characteristics could explain heterogeneity in effect sizes. Attrition in the cohort did not clearly moderate any estimate (*p* > .05). Methodological quality (*p* = .0072) and sample size (*p* = .0225) were negatively associated with the magnitude of genetic nurture effects, which can be attributed to the potentially influential study (Kong et al., 2018) with the highest quality score and sample size (see sTables 7 in the supplement).

After adjusting for phenotypic family-level factors (i.e., parental educational level or family SES), genetic nurture effects attenuated to a large extent (kunadjusted = 22, *β*_unadjusted_ = 0.07, 95% CI [0.07, 0.08] vs. kadjusted = 18, *β*_adjusted_ = 0.02, 95% CI [0.01, 0.03], *p*_adjustment_ < .0001), for more details see section 6 in the supplement.

## Discussion

Across 12 studies including 38,654 distinct parent(s)-offspring pairs or trios from eight cohorts, we found strong evidence for genetic nurture, i.e., parental genotypes affecting child outcomes via environmental pathways. The magnitude of genetic nurture effects is similar in both parents and is about half the size of direct genetic effects originating in the offspring due to genetic inheritance. In the following sections, we discuss in turn: (a) the impact of genetic nurture and direct genetic effects on offspring’s educational outcome, (b) sources of heterogeneity in genetic impacts, and (c) implications.

### Genomic Prediction of Education: Evidence for Genetic Nurture and Direct Genetic Effects

Pooled estimates showed significant effects of parental genetic propensity for educational attainment on offspring educational outcomes. Importantly, effects of parental genetic propensity for education on offspring’s educational outcomes can reflect both genetic nurture and direct genetic transmission. We observed a small effect of genetic nurture (*β*_genetic nurture_ = 0.08) on educational outcomes, which is similar to estimates obtained from family-informed designs not including measured genomic data, such as adoption designs (*β* = 0.10)(Domingue & Fletcher, 2020).

Relative to effects of genetic nurture, we observed larger direct genetic effects on educational outcomes. Our pooled estimate of the unadjusted effect of offspring genotype (*β*_child unadjusted_ = 0.21) is comparable to estimates obtained from studies assessing the explanatory power of EA PGS on one’s educational outcome without accounting for genetic nurture effects, which typically range between *β* = 0.15 and *β* = 0.39 (Allegrini et al., 2019; Belsky et al., 2016; Domingue et al., 2015; von Stumm et al., 2020). Direct genetic effects, which correspond to effects of the offspring’s genotype on their own phenotype, are free from inflation due to genetic nurture. Our pooled estimate of direct genetic effects (*β*_direct genetic_ = 0.17) corresponds to the lower bound of previous genomic predictions of educational outcomes within twin pairs (e.g., *β* = 0.17-27 (Selzam et al., 2019; Willoughby et al., 2019).

Genetic nurture effects are about half the size of direct genetic effects (*β*_genetic nurture_ */β*_direct genetic_ = 0.47) in our pooled estimate, which is consistent with the largest study of genetic nurture effects so far (ratio = 0.43) even when this potentially influential study (Kong et al., 2018)^1^ is excluded from the meta-analysis. Recent studies have also implemented other methods to investigate genetic nurture effects with twin (Selzam et al., 2019) and adoption (Cheesman et al., 2020; Domingue & Fletcher, 2020) designs. As evidence from these alternative designs accumulate, it will be key to examine the consistency of estimates across designs to strengthen confidence in findings on genetic nurture for educational outcomes (Lawlor, Tilling, & Davey Smith, 2017).

It is worth noting that the evidence included in this meta-analysis can only detect genetic nurture and direct genetic effects to the extent that the PGS capture heritability in educational outcomes. To date, PGS based on the most accurate GWASs still only capture a fraction of heritability (Allegrini et al., 2019; Cesarini & Visscher, 2017). A recent study using Relatedness Disequilibrium Regression (RDR), which estimated heritability by exploiting variation in relatedness due to random Mendelian segregation, provided a ‘ceiling’ for potential gain from increasing the predictive accuracy of PGS (Young et al., 2018). We found our estimate of genetic nurture based on PGS explained 0.64% (*β*_genetic nuture_^2^) of child educational outcomes versus 6.6% for RDR, and direct genetic effects based on PGS explained 2.89% (*β*_direct genetic_^2^) versus 17% for RDR (Young et al., 2018).

While missing heritability may lead to underestimating the true extent of genetic nurture, assortative mating and population stratification may have inflated our genetic nurture effects (Shen & Feldman, 2020). Bias resulting from assortative mating is thought to be small in magnitude (Kong et al., 2018; Shen & Feldman, 2020). Population stratification was controlled for by using principal component analysis in most studies included in the meta-analysis but residual population stratification may still exist. Emerging methods should, in future, better account for remaining biases for example by capitalizing further on family-based designs (Balbona, Kim, & Keller, 2020; Demange et al., 2020; Young et al., 2020).

### Genomic Prediction of Education: Sources of Heterogeneity

First, when comparing estimates of genomic measures from different parents, we found similar genetic nurture effects regardless of parent of origin. One explanation of such findings is that both parents are equally as important in shaping the environment that, in turn, influences their offspring’s educational outcomes. Our findings do not preclude that parents may contribute by different mechanisms (e.g. via distal factors like increased family income or by proximal factors like reading to the child). Behavioural studies showed that the relationship between parental involvement and children’s educational achievement was equally strong for fathers and mothers (Barger et al., 2019; Kim & Hill, 2015). A renewed emphasis on the role of fathers is needed and, whenever possible, fathers should systematically be included in research and intervention efforts. Research in the area should turn to examining more systematically genetic nurture effects in alternative family arrangements (e.g., single-parent families) and according to varying level of parental involvement (e.g., separated families where one parent is mostly uninvolved). We expect genetic nurture effects to vary accordingly, e.g., be stronger for the most involved parent, which should shed further light on environmental factors mediating genetic nurture effects. Another explanation of similar effect sizes for both parents could be that genetic nurture operates through the broad family-level environment which is shared by both parents (e.g., school quality in the neighbourhood). Future investigations are required to identify such environmental mediators.

Second, the magnitude of genetic nurture effects was slightly smaller in the virtual parent design versus the statistical control approach. A recent study has shown the importance of using complete trio data, as missing the genotype of one parent can bias direct genetic effects and genetic nurture effects (Tubbs, Zhang, & Sham, 2020). Evidence from one of the included studies (T. Morris et al., 2020) using both partial and full statistical control approach echoed this view. In our meta-analysis however, we found no strong evidence for differences based upon findings from partial control (one parent) or full control (two parents). This may however reflect that these controls were estimated from different samples, hiding true differences. Additional work is needed to compare estimates of genetic nurture using different analytical methods within the same sample to better understand the equivalence and comparability of different approaches.

Third, we expected to find larger genetic nurture effects on educational attainment relative to educational achievement, since the genomic measures were constructed using GWAS of educational attainment. This was verified when restricting the analysis to studies where both educational attainment and achievement were assessed (de Zeeuw et al., 2020; Rustichini, Iacono, Lee, & McGue, 2018). Another explanation of larger genetic nurture effects for educational attainment compared to educational achievement is that attainment may be more socially influenced than achievement (T. T. Morris, Davies, & Smith, 2020). That is, may be easier for parents to influence attainment (e.g. by accessing more exclusive schooling or financially supporting further education). However, developmental trends in genetic nurture effects warrant more investigation. Heritability of educational outcomes increases with age (T. Morris, Davies, Dorling, Richmond, & Smith, 2018). Conversely, the nurturing behaviours from parents may impact offspring more at earlier ages, as they spend more time at home rather than school, spend more time with their parents rather than peers, which might lead to genetic nurture effects decreasing with age. Moreover, genetic nurture may act distinctively over time through different pathways as suggested by a recent study showing parent non-cognitive but not cognitive related characteristics were more important for educational achievement at age 16 than age 12 (Demange et al., 2020).

For direct genetic effects, estimates for educational achievement were significantly larger than attainment. This finding agrees with previous twin evidence, which suggested ~60% heritability for educational achievement measured in childhood and adolescence (Rimfeld et al., 2018) and ~40% for educational attainment measured in adulthood (Branigan, McCallum, & Freese, 2013; Silventoinen et al., 2020). Some plausible explanations of such higher heritability/direct genetic effects for educational achievement are discussed in the supplement (section 7.5).

Fourth and as expected and consistent with previous studies (e.g., Bates et al., 2019; Selzam et al., 2017), the predictive accuracy of the GWAS used to construct the PGS significantly moderated effect sizes of genetic nurture, with more accurate GWAS associated with larger effect estimates.

Methodological quality and sample size were negatively associated with genetic nurture effects in a modest manner, which may reflect the impact of the largest and best quality study included (Kong et al., 2018). Nevertheless, it suggests that more reliable studies, namely with more rigorous methodology and larger sample size, may produce more conservative estimates of genetic nurture effects.

Lastly, we found that accounting for observed measures of parental education or family SES decreased the effect of genetic nurture by ¾. This suggests that a substantial amount of genetic nurture effects can be attributed to environmental pathways directly related to parental education, occupation and income. It echoes the view that offspring’s educational outcomes are influenced by the availability of resources in their family, either indicated by socioeconomic background or the education of their parents (Björklund & Salvanes, 2011; Shavit & Blossfeld, 1993; Sirin, 2005). The finding that genetic nurture operates, to a large extent, on broad family-level environment helps to explain why both parents may have similar genetic nurture effects on offspring educational outcomes. Future investigations should explore specific family-level pathways through which genetic nurture operate to inform compensatory interventions (e.g., financial support vs. schooling access). Importantly, findings that broad family-level social economic characteristics largely explain genetic nurture effects do not preclude the importance of proximal factors such as parenting in the chain of factors leading to educational outcomes.

### Implications

Genetic nurture had a robust effect on children’s educational outcomes, similar across parents. As genetic nurture effects are free from the genetic confounding induced by the genetic relatedness between parents and offspring, it suggests that the environment created by parents impacts on offspring educational outcomes independent of genetic transmission. Although the magnitude of such an effect is small when interpreting on conventional metrics (Cohen, 1988), it is important to note it will likely increase as the explanatory power of polygenic scores increases. In addition, such an effect size is proposed to be of medium policy importance on education interventions (Kraft, 2020). More thorough investigations of specific environmental pathways through which genetic nurture operate may help in understanding how educational outcomes are maintained across generations and help design better compensatory interventions. Such interventions could target environmental pathways in two ways, either targeting distal risk factors for educational outcomes (e.g. parental education, income distribution, equal access to good quality schooling) or more proximal pathways (rearing environment such as parenting). Such interventions can, to some extent, be expected to disrupt the intergenerational cycle of educational underachievement and foster social mobility. It is important to note that how well children do in school does depend to a substantial degree on the genetic lottery (i.e., inheriting more genetic variants associated with educational success), which policy makers often overlook (Dewar et al., 2017). At a broader level, our findings provide among the strongest evidence to date that differences in education are consistently influenced by endogenous sources of educational inequalities (e.g. one’s own genetics) and exogenous sources of inequalities including genetic nurture effects originating in parents and mediated partially through broad-level family characteristics like SES. All these endogenous and exogenous sources of educational inequalities are beyond individual responsibility/control and therefore may be construed as justification for compensatory interventions. Interventions could thus aim to provide more support to children in need throughout their education, contributing to a building a fairer society with more equitable educational opportunities.

### Limitations

We cannot completely rule out bias from unmeasured assortative mating, residual population stratification and sibling genetic nurture (Demange et al., 2020; Young et al., 2020), which may inflate genetic nurture effects. In addition, all included studies were based on European ancestry populations and thus have a profound Eurocentric bias. The generalizability of our estimates to non-European population is unclear as genomic measures are not necessarily accurate across populations (Martin et al., 2019). For example, PGS constructed from EA3, which was conducted in white Europeans, captures 10.6% of the variation of educational attainment in white Americans but only about 1.6% of the variation among African Americans (Lee et al., 2018). Third, differential measurement error in outcomes may affect genetic (nurture) effect sizes. Comparison between different outcome types (e.g. attainment and achievement) should therefore be interpreted with caution.

## Acknowledgments

B.W. and JB.P. are funded by a Nuffield Foundation project (EDO/43939); J.R.B is funded by a Wellcome Trust Sir Henry Wellcome fellowship (215917/Z/19/Z); T.S. is funded by a Wellcome Trust Sir Henry Wellcome fellowship (218641/Z/19/Z). D.B. is funded by the Economic and Social Research Council (ES/M001660/1) and Medical Research Council (MR/V002147/1). F.D. is a consultant for University College London funded by the Nuffield Foundation project.

## Footnotes

1 In Kong et al. 2018, estimated genetic nurture effects were 0.067, estimated direct effects were 0.157. The ratio of genetic nurture/direct genetic effects is thus 0.067/0.157 = 0.43.

## References

Allegrini, A. G., Selzam, S., Rimfeld, K., von Stumm, S., Pingault, J.-B., & Plomin, R. (2019). Genomic prediction of cognitive traits in childhood and adolescence. Molecular Psychiatry, 24(6), 819–827.

Armstrong-Carter, E., Trejo, S., Hill, L. J. B., Crossley, K. L., Mason, D., & Domingue, B. W. (2020). The Earliest Origins of Genetic Nurture: The Prenatal Environment Mediates the Association Between Maternal Genetics and Child Development. Psychological science. Retrieved from <Go to ISI>://WOS:000537515200001

Assink, M., & Wibbelink, C. J. (2016). Fitting three-level meta-analytic models in R: A step-by-step tutorial. The Quantitative Methods for Psychology, 12(3), 154–174.

Ayorech, Z., Krapohl, E., Plomin, R., & von Stumm, S. (2017). Genetic influence on intergenerational educational attainment. Psychological science, 28(9), 1302–1310.

Balbona, J., Kim, Y., & Keller, M. C. (2020). Estimation of parental effects using polygenic scores. bioRxiv, 2020.2008.2011.247049. doi:10.1101/2020.08.11.247049

Barger, M. M., Kim, E. M., Kuncel, N. R., & Pomerantz, E. M. (2019). The relation between parents’ involvement in children’s schooling and children’s adjustment: A meta-analysis. Psychological bulletin, 145(9), 855–890. doi:10.1037/bul0000201

Bates, T. C., Maher, B. S., Colodro-Conde, L., Medland, S. E., McAloney, K., Wright, M. J., . . . Gillespie, N. A. (2019). Social competence in parents increases children’s educational attainment: Replicable genetically-mediated effects of parenting revealed by non-transmitted DNA. Twin Research and Human Genetics, 22(1), 70–74. Retrieved from http://journals.cambridge.org/action/displayJournal?jid=THG http://ovidsp.ovid.com/ovidweb.cgi?T=JS&PAGE=reference&D=emexa&NEWS=N&AN=626076119

Bates, T. C., Maher, B. S., Medland, S. E., McAloney, K., Wright, M. J., Hansell, N. K.,…Gillespie, N. A. (2018). The Nature of Nurture: Using a Virtual-Parent Design to Test Parenting Effects on Children’s Educational Attainment in Genotyped Families. Twin Research and Human Genetics, 21(2), 73–83. Retrieved from http://journals.cambridge.org/action/displayJournal?jid=THG http://ovidsp.ovid.com/ovidweb.cgi?T=JS&PAGE=reference&D=emed19&NEWS=N&AN=621226923

Belsky, D. W., Moffitt, T. E., Corcoran, D. L., Domingue, B., Harrington, H., Hogan, S.,…Williams, B. S. (2016). The genetics of success: How single-nucleotide polymorphisms associated with educational attainment relate to life-course development. Psychological science, 27(7), 957–972.

Björklund, A., & Salvanes, K. G. (2011). Chapter 3 - Education and Family Background: Mechanisms and Policies. In E. A. Hanushek, S. Machin, & L. Woessmann (Eds.), Handbook of the Economics of Education (Vol. 3, pp. 201–247): Elsevier.

Branigan, A. R., McCallum, K. J., & Freese, J. (2013). Variation in the heritability of educational attainment: An international meta-analysis. Social Forces, 92(1), 109–140.

Cesarini, D., & Visscher, P. M. (2017). Genetics and educational attainment. NPJ science of learning, 2(1), 1–7.

Cheesman, R., Hunjan, A., Coleman, J. R., Ahmadzadeh, Y., Plomin, R., McAdams, T. A.,…Breen, G. (2020). Comparison of adopted and nonadopted individuals reveals gene–environment interplay for education in the UK Biobank. Psychological science, 31(5), 582–591.

Cohen, J. (1988). Statistical Power Analysis for the Behavioral Sciences, (L. Erlbaum Associates, Hillsdale, NJ). In: Erlbaum Associates: Hillsdale, NJ, USA.

Conti, G., Heckman, J., & Urzua, S. (2010). The education-health gradient. American Economic Review, 100(2), 234–238.

Crespo, L., López-Noval, B., & Mira, P. (2014). Compulsory schooling, education, depression and memory: New evidence from SHARELIFE. Economics of Education Review, 43, 36–46.

de Zeeuw, E. L., Hottenga, J. J., Ouwens, K. G., Dolan, C. V., Ehli, E. A., Davies, G. E.,…van Bergen, E. (2020). Intergenerational Transmission of Education and ADHD: Effects of Parental Genotypes. Behavior Genetics. doi:10.1007/s10519-020-09992-w

Del Re, A. C. (2013). compute.es: Compute Effect Sizes. Retrieved from https://cran.r-project.org/package=compute.es

Demange, P. A., Hottenga, J. J., Abdellaoui, A., Malanchini, M., Domingue, B. W., de Zeeuw, E. L.,…van Bergen, E. (2020). Parental influences on offspring education: indirect genetic effects of non-cognitive skills. bioRxiv.

Dewar, M., Esser, F. C., Benczur, P., Campolongo, F., Harasztosi, P., Karagiannis, S.,…Ivaskaite-Tamosiune, V. (2017). What makes a fair society? Insights and evidence. Retrieved from

Domingue, B. W., Belsky, D. W., Conley, D., Harris, K. M., & Boardman, J. D. (2015). Polygenic influence on educational attainment: New evidence from the National Longitudinal Study of Adolescent to Adult Health. Aera Open, 1(3), 2332858415599972.

Domingue, B. W., & Fletcher, J. (2020). Separating measured genetic and environmental effects: Evidence linking parental genotype and adopted child outcomes. Behavior Genetics, 1–9.

Dubow, E. F., Boxer, P., & Huesmann, L. R. (2009). Long-term effects of parents’ education on children’s educational and occupational success: Mediation by family interactions, child aggression, and teenage aspirations. Merrill-Palmer quarterly (Wayne State University. Press), 55(3), 224.

Dudbridge, F. (2013). Power and predictive accuracy of polygenic risk scores. PLoS Genet, 9(3), e1003348.

Foundation, R. S., Kaplan, G. A., House, J. S., Schoeni, R. F., & Pollack, H. A. (2008). Making Americans healthier: Social and economic policy as health policy: Russell Sage Foundation.

Hertz, T., Jayasundera, T., Piraino, P., Selcuk, S., Smith, N., & Verashchagina, A. (2008). The inheritance of educational inequality: International comparisons and fifty-year trends. The BE Journal of Economic Analysis & Policy, 7(2).

Kim, S. w., & Hill, N. E. (2015). Including fathers in the picture: A meta-analysis of parental involvement and students’ academic achievement. Journal of Educational Psychology, 107(4), 919–934. doi:10.1037/edu0000023

Koellinger, P. D., & Harden, K. P. (2018). Using nature to understand nurture. Science, 359(6374), 386–387.

Kong, A., Thorleifsson, G., Frigge, M. L., Vilhjalmsson, B. J., Young, A. I., Thorgeirsson, T. E.,…Masson, G. (2018). The nature of nurture: Effects of parental genotypes. Science, 359(6374), 424–428.

Kraft, M. A. (2020). Interpreting effect sizes of education interventions. Educational Researcher, 49(4), 241–253.

Krapohl, E., & Plomin, R. (2016). Genetic link between family socioeconomic status and children’s educational achievement estimated from genome-wide SNPs. Molecular Psychiatry, 21(3), 437–443.

Lawlor, D. A., Tilling, K., & Davey Smith, G. (2017). Triangulation in aetiological epidemiology. International Journal of Epidemiology, 45(6), 1866–1886. doi:10.1093/ije/dyw314

Lee, J. J., Wedow, R., Okbay, A., Kong, E., Maghzian, O., Zacher, M.,…Linnér, R. K. (2018). Gene discovery and polygenic prediction from a 1.1-million-person GWAS of educational attainment. Nature Genetics, 50(8), 1112.

Martin, A. R., Kanai, M., Kamatani, Y., Okada, Y., Neale, B. M., & Daly, M. J. (2019). Clinical use of current polygenic risk scores may exacerbate health disparities. Nature Genetics, 51(4), 584–591. doi:10.1038/s41588-019-0379-x

Moher, D., Liberati, A., Tetzlaff, J., Altman, D. G., & Group, P. (2009). Preferred reporting items for systematic reviews and meta-analyses: the PRISMA statement. PLoS medicine, 6(7), e1000097.

Morris, T., Davies, N., Dorling, D., Richmond, R., & Smith, G. D. (2018). Testing the validity of value-added measures of educational progress with genetic data. British Educational Research Journal, 44(5), 725–747. doi:https://doi.org/10.1002/berj.3466

Morris, T., Davies, N., Hemani, G., & Smith, G. D. (2020). Population phenomena inflate genetic associations of complex social traits. Science Advances, 6(16). Retrieved from <Go to ISI>://WOS:000528276800007

Morris, T. T., Davies, N. M., & Smith, G. D. (2020). Can education be personalised using pupils’ genetic data? eLife, 9, e49962.

Okbay, A., Beauchamp, J. P., Fontana, M. A., Lee, J. J., Pers, T. H., Rietveld, C. A.,…Meddens, S. F. W. (2016). Genome-wide association study identifies 74 loci associated with educational attainment. Nature, 533(7604), 539–542.

R Core Team. (2019). R: A language and environment for statistical computing. Vienna, Austria: R Foundation for Statistical Computing.

Rietveld, C. A., Medland, S. E., Derringer, J., Yang, J., Esko, T., Martin, N. W.,…Agrawal, A. (2013). GWAS of 126,559 individuals identifies genetic variants associated with educational attainment. Science, 340(6139), 1467–1471.

Rimfeld, K., Malanchini, M., Krapohl, E., Hannigan, L. J., Dale, P. S., & Plomin, R. (2018). The stability of educational achievement across school years is largely explained by genetic factors. NPJ science of learning, 3(1), 1–10.

Rodgers, M. A., & Pustejovsky, J. E. (2020). Evaluating meta-analytic methods to detect selective reporting in the presence of dependent effect sizes. Psychological Methods.

Ronald, A. (2020). Polygenic scores in child and adolescent psychiatry–strengths, weaknesses, opportunities and threats. Journal of Child Psychology and Psychiatry, 61(5), 519–521.

Rustichini, A., Iacono, W. G., Lee, J., & McGue, M. (2018). Polygenic score analysis of educational achievement and intergenerational mobility.

Selzam, S., Krapohl, E., von Stumm, S., O’Reilly, P., Rimfeld, K., Kovas, Y.,…Plomin, R. (2017). Predicting educational achievement from DNA. Molecular Psychiatry, 22(2), 267–272.

Selzam, S., Ritchie, S. J., Pingault, J.-B., Reynolds, C. A., O’Reilly, P. F., & Plomin, R. (2019). Comparing within-and between-family polygenic score prediction. The American Journal of Human Genetics, 105(2), 351–363.

Shavit, Y., & Blossfeld, H.-P. (1993). Persistent Inequality: Changing Educational Attainment in Thirteen Countries. Social Inequality Series: ERIC.

Shen, H., & Feldman, M. W. (2020). Genetic nurturing, missing heritability, and causal analysis in genetic statistics. Proceedings of the national academy of sciences, 117(41), 25646–25654.

Silventoinen, K., Jelenkovic, A., Sund, R., Latvala, A., Honda, C., Inui, F.,…Kaprio, J. (2020). Genetic and environmental variation in educational attainment: an individual-based analysis of 28 twin cohorts. Scientific reports, 10(1), 12681. doi:10.1038/s41598-020-69526-6

Sirin, S. R. (2005). Socioeconomic status and academic achievement: A meta-analytic review of research. Review of Educational Research, 75(3), 417–453.

Sorensen, H. J., Debost, J. C., Agerbo, E., Benros, M. E., McGrath, J. J., Mortensen, P. B.,…Petersen, L. Polygenic Risk Scores, School Achievement, and Risk for Schizophrenia: A Danish Population-Based Study. Biological psychiatry, 84(9), 684–691. Retrieved from <Go to ISI>://WOS:000446411600012

Stang, A. (2010). Critical evaluation of the Newcastle-Ottawa scale for the assessment of the quality of nonrandomized studies in meta-analyses. European journal of epidemiology, 25(9), 603–605.

Stroup, D. F., Berlin, J. A., Morton, S. C., Olkin, I., Williamson, G. D., Rennie, D.,…Thacker, S. B. (2000). Meta-analysis of observational studies in epidemiology: a proposal for reporting. Jama, 283(15), 2008–2012.

Tubbs, J. D., Zhang, Y. D., & Sham, P. C. (2020). Intermediate confounding in trio relationships: The importance of complete data in effect size estimation. Genetic Epidemiology, 44(4), 395–399.

Viechtbauer, W. (2010). Conducting meta-analyses in R with the metafor package. Journal of statistical software, 36(3), 1–48.

Visscher, P. M., Wray, N. R., Zhang, Q., Sklar, P., McCarthy, M. I., Brown, M. A., & Yang, J. (2017). 10 years of GWAS discovery: biology, function, and translation. The American Journal of Human Genetics, 101(1), 5–22.

von Stumm, S., Smith-Woolley, E., Ayorech, Z., McMillan, A., Rimfeld, K., Dale, P. S., & Plomin, R. (2020). Predicting educational achievement from genomic measures and socioeconomic status. Developmental science, 23(3), e12925.

Willoughby, E. A., McGue, M., Iacono, W. G., Rustichini, A., & Lee, J. J. (2019). The role of parental genotype in predicting offspring years of education: evidence for genetic nurture. Molecular Psychiatry. Retrieved from http://www.nature.com/mp/index.html http://ovidsp.ovid.com/ovidweb.cgi?T=JS&PAGE=reference&D=emexa&NEWS=N&AN=2002635365

Young, A. I., Frigge, M. L., Gudbjartsson, D. F., Thorleifsson, G., Bjornsdottir, G., Sulem, P.,…Kong, A. (2018). Relatedness disequilibrium regression estimates heritability without environmental bias. Nature Genetics, 50(9), 1304–1310. doi:10.1038/s41588-018-0178-9

Young, A. I., Nehzati, S. M., Lee, C., Benonisdottir, S., Cesarini, D., Benjamin, D. J.,…Kong, A. (2020). Mendelian imputation of parental genotypes for genome-wide estimation of direct and indirect genetic effects. bioRxiv.

