## Supplementary Notes and Figures for "Genetic nurture effects on education: a systematic review and meta-analysis"

### SUPPLEMENTARY MATERIAL

#### Table of Contents

1. CAPTURING GENETIC NURTURE EFFECTS WITH PARENT(S)-OFFSPRING GENOTYPE 4

**1.1 Virtual-parent design 4**

**1.2 Statistical control approach 5**

2. GENOME-WIDE ASSOCIATION STUDIES AND POLYGENIC SCORES OF EDUCATIONAL ATTAINMENT 6

3. STUDY SELECTION AND ASSESSMENT 7

**3.1 Literature search 7**

**3.2 Inclusion criteria 9**

**3.3 Assessment of methodological quality 10**

4. DATA EXTRACTION AND SYNTHESIS 13

**4.1 Effect size calculation 13**

**4.2 Multilevel Random Effects Model (MREM) 14**

**4.3 Moderator analyses of study characteristics 15**

5. UNADJUSTED PARENTAL AND CHILD EFFECTS 15

**5.1 Unadjusted parental effects 15**

**5.2 Unadjusted child effects 16**

6. FAMILY-LEVEL ADJUSTMENT 17

7. SENSITIVITY ANALYSES 18

**7.1 Robust confidence intervals of dependent estimates 18**

**7.2 Impact of recalibrating estimates using the average parental PGS 19**

**7.3 Impact of a potentially influential study 20**

**7.4 Jack-knife leave-one-out analyses 21**

**7.5 The moderating effect of outcome type within study 21**

8. SFIGURES 23

**8.1 sFigure 1. Funnel plots for parental effects on educational outcomes 23**

**8.2 sFigure 2. Jackknife sensitivity analyses for parental effects on educational outcomes 24**

**8.3 sFigure 3. Funnel plots for child effects on educational outcomes 25**

**8.4 sFigure 4. Jackknife sensitivity analyses for child effects on educational outcomes 26**

9. SREFERENCES 27

### 1. Capturing genetic nurture effects with parent(s)-offspring genotype

#### 1.1 Virtual-parent design

The parental genetic material transmitted to their offspring is randomly assigned during meiosis, with each allele having a 50% chance of being transmitted to the gamete (egg or sperm) and then to the offspring. Alleles that are not transmitted to offspring can nonetheless influence offspring outcomes through environmental rather than genetic pathways. The non-transmitted parental genotype can be considered as a "virtual parent" who is not genetically related to the offspring, whereas the transmitted parental genotype (50% from mother and 50% from father) makes up the child genotype. The effect of polygenic scores (PGS) derived from non-transmitted parental genotype on offspring outcome is thus free from genetic confounding between parents and offspring due to shared genotype.

In regressions with child education as the outcome, let $PGS_T$ and $PGS_{NT}$ be the standardized PGS of transmitted and non-transmitted parental genotype respectively, Cp be the child phenotype, and $\beta_T$ and $\beta_{NT}$ be the corresponding respective coefficients. The model can be expressed as follows:

$$Cp = \beta_T * PGS_T + \beta_{NT} * PGS_{NT} + e$$

The estimated genetic nurture effects are:

$$\text{Genetic nurture effects} = \beta_{NT}$$

As both transmitted and non-transmitted parental genotype have nurturing effects, direct genetic effects originating in the child are:

$$\text{Direct genetic effects} = \beta_T - \beta_{NT}$$

For detailed decompositions of aforementioned equations see (Shen & Feldman, 2020). For more sophisticated decompositions of genetic influences using the virtual parent design, including genetic nurture effects, direct genetic effects, the assortative mating–induced confounding effect for the direct genetic effect component, and the confounding effect of the genetic nurturing component, see (Kong et al., 2018).

##### 1.2 Statistical control approach

The statistical control approach utilizes the complete parental genotype as an aggregation of transmitted and non-transmitted genotypes. Disentanglement of genetic nurture and direct genetic effects is achieved by modelling the association between parental PGS and offspring phenotype while controlling for offspring PGS. As offspring genotype can fully mediate the effects of the parental transmitted genotype, the remaining effects of parental genotype on offspring outcome can only be environmentally mediated, i.e., via genetic nurture effects. Similarly, since the effect of offspring genotype is controlled for parental genotype, it thus reflects direct genetic effects free from the inflation from genetic nurture.

Let $PGS_C$, $PGS_M$ and $PGS_P$ be the standardized PGS of child, maternal and paternal genotype respectively, and $\beta_C$, $\beta_M$ and $\beta_P$ be the corresponding coefficients. The full statistical control model can be expressed as follows when genotypes of both parents are available:

$$Cp = \beta_C * PGS_C + \beta_M * PGS_M + \beta_P * PGS_P + e$$

The partial statistical control model can be expressed as follows when genotypes of one parent is available:

$$Cp = \beta_C * PGS_C + \beta_M * PGS_M + e$$

or

$$Cp = \beta_C * PGS_C \beta_P * PGS_P + e$$

The estimated genetic nurture effects are:

$$\text{Maternal genetic nurture effects} = \beta_M$$

$$\text{Paternal genetic nurture effects} = \beta_P$$

It is important to note, assortative mating, i.e., the phenotypic correlation between the mother and the father, could still bias the estimation where the genotype of only one parent is available.

The estimated direct genetic effects are:

$$\text{Direct genetic effects} = \beta_C$$

#### 2. Genome-wide association studies and polygenic scores of educational attainment

The first EA GWAS (EA1) was conducted in 2013 in a discovery sample of 101,069 individuals, and found three independent SNPs with genome-wide significance (i.e., *p* value threshold of $5 \times 10^{-8}$)(Rietveld et al., 2013). The discovery sample was extended to 293,723 individuals in 2016 by the second EA GWAS (EA2), which identified 74 genome-wide significant loci associated with years of schooling completed (Okbay et al., 2016). The most recent EA GWAS (EA3) was conducted in 2018 in a sample of approximately 1.1 million individuals ($N$ = 1,131,881), the marked increase in sample size boosted the predictive accuracy/statistical power to detect genetic associations and resulted in identifying 1,271 independent genome-wide-significant SNPs (Lee et al., 2018). However, individual SNP effect sizes are extremely small (Chabris, Lee, Cesarini, Benjamin, & Laibson, 2015; Gratten, Wray, Keller, & Visscher, 2014), and using alleles with genome-wide significance only explain a trivial amount of the total variance in outcomes. In response, researchers explored the idea of relaxing the significance threshold and summing these individual effects together to give an overall estimate of genetic liability for a trait - a polygenic score (PGS) (Dudbridge, 2013; Ronald, 2020). PGS are calculated by summing the effects of many common genetic variants, which are weighted by effect sizes derived from corresponding GWAS results (International Schizophrenia Consortium, 2009; Wray, Goddard, & Visscher,

2007). PGS from all measured SNPs accounted for about 2% of the variance in years of schooling using EA1, and 3.2% and 11% using EA2 and EA3, respectively.

#### 3. Study selection and assessment

##### 3.1 Literature search

Two search strategies were employed for study identification. First, we systematically searched the following databases:

1) Ovid:

i) MEDLINE In-Process & Other Non-Indexed Citations and Daily

ii) EMBASE

iii) PsycINFO

2) Web of Science Core Collection:

i) Sciences Citation Index Expanded (SCI-EXPANDED);

ii) Social Sciences Citation Index (SSCI);

iii) Arts & Humanities Citation Index (A&HCI);

iv) Emerging Sources Citation Index (ESCI).

3) PubMed.

The search terms are described below:

((education* [Title/Abstract]) OR (academic [Title/Abstract]) OR (college degree [Title/Abstract]) OR (college entry [Title/Abstract]) OR (university degree [Title/Abstract]) OR (university entry [Title/Abstract]) OR (school performance* [Title/Abstract]) OR (school achievement* [Title/Abstract]) OR (performance in school [Title/Abstract]) OR (schooling [Title/Abstract])

AND

(polygenic scor* [Title/Abstract]) OR (polygenic risk scor* [Title/Abstract]) OR (genetic scor* [Title/Abstract]) OR (genetic risk scor* [Title/Abstract]) OR (genomic scor* [Title/Abstract]) OR (genomic risk scor* [Title/Abstract]) OR (genome-wide polygenic scor* [Title/Abstract])

AND

(nature of nurture [Title/Abstract]) OR (genetic nurtur* [Title/Abstract]) OR (virtual-parent design [Title/Abstract]) OR (pseudo-control [Title/Abstract]) OR (dynastic effect* [Title/Abstract]) OR (intergeneration* [Title/Abstract]) OR (multigeneration* [Title/Abstract]) OR (transmit* allele* [Title/Abstract]) OR (nontransmit* allele* [Title/Abstract]) OR (non-transmit* allele* [Title/Abstract]) OR (passive gene–environment correlation [Title/Abstract]) OR (genetic inheritance [Title/Abstract]) OR (genetic confounding* [Title/Abstract]) OR (genetic transmission [Title/Abstract]) OR (genetic influence* [Title/Abstract]) OR (parental transmission [Title/Abstract]) OR (maternal transmission [Title/Abstract]) OR (paternal transmission [Title/Abstract]) OR (parental influence* [Title/Abstract]) OR (maternal influence* [Title/Abstract]) OR (paternal influence* [Title/Abstract]) OR (social inheritance [Title/Abstract]) OR (social genetic effect* [Title/Abstract]) OR (environmental transmission [Title/Abstract]) OR (cultural transmission [Title/Abstract]) OR (vertical transmission [Title/Abstract]) OR (familial transmission [Title/Abstract]) OR (family-based stud* [Title/Abstract]) OR (trio* [Title/Abstract] OR (triad* [Title/Abstract] OR (dual* [Title/Abstract] OR (dyad* [Title/Abstract]))

These search terms were translated into suitable terms for the Ovid database (MEDLINE, EMBASE and PsycINFO), Web of Science and PubMed, which can be made available upon request.

Second, we manually searched the reference list of relevant articles, including studies that used virtual parent and statistical control approaches to investigate the genetic nurture effects on education, as well as unpublished evidence in preprint platforms were screened to identify articles that were missed by the search. Two authors (B.W. and T.S.) independently screened all articles retrieved from the search on Rayyan (http://rayyan.qcri.org), a free platform for systematic review (Ouzzani, Hammady, Fedorowicz, & Elmagarmid, 2016). Potentially eligible studies (see criteria below) were then reviewed in full text. This search resulted in n=12 studies eligible for inclusion.

#### 3.2 Inclusion criteria

Studies were included if they assessed genetic nurture effects on educational outcomes, either educational attainment (e.g., years of education completed, highest degree obtained) or educational achievement (e.g., national tests scores or levels, school grades) in the general population. Due to lack of data in preliminary searches, it was part of the protocol to exclude studies that were conducted in clinically-referred sample or focused exclusively on specific types of educational-related outcomes (e.g., performances on math course or linguistics). No genetic nurture study on clinical sample or specific educational outcomes was present after the systematic search. Studies were required to use one of the two designs that rely on genotype data from parents and their biological offspring: 1) virtual parent: testing genetic nurture effects on education by using parent(s)' non-transmitted genotype to predict children's education. In this case polygenic scores of educational attainment (hereafter referred as EA PGS) calculated from at least one parent's non-transmitted alleles should be used; 2) statistical control: testing genetic nurture effects on education by using parent(s)' whole genotype to predict children's education over and above children's own genotype, in this case the EA PGS of child and at least one parent should be used. Twin evidence relying on siblings (Cheesman et al., 2020; Selzam et al., 2019) and adoption evidence relying on adoption relations (Domingue & Fletcher, 2020) were not included in the meta-analysis to avoid introducing additional heterogeneity but served as measure of comparability and robustness when discussing results of our pooled genetic nurture effects.

##### 3.3 Assessment of methodological quality

An adapted version of the Newcastle - Ottawa Quality Assessment Scale for Cohort Studies (Stang, 2010) was applied to evaluate the methodological quality of the included studies. The following scoring criteria was used:

***Methodological quality assessment criteria of studies on genetic nurture effects on education***

Note: A study can be awarded a maximum of one point for each numbered item within all categories.

Representativeness and attrition

1) Population representativeness of the cohort

    a) truly representative of the average cohort in the community, e.g. pregnancy/birth cohort, epidemiological cohort, population-based (1)

    b) somewhat representative of the average cohort in the community, e.g., twin cohort, family-based (0.5)

    c) selected group of users, e.g., case-control, genomics company (0)

    d) no description of the derivation of the cohort (0)

2) Attrition in the cohort (due to genotyping or outcome availability)

    a) complete cohort - all subjects of original cohort are used (1)

    b) subjects lost unlikely to introduce bias - small number lost - > 70 % of original cohort are used, or description provided of those lost[1] (1)

c) < 70% of original cohort are used and no description of those lost (0)

d) no statement (0)

[1]only code 1 when subjects lost is unlikely to introduce bias, e.g., if description indicate random lost.

Exposure: polygenic scores (PGS) of educational attainment

3) Power/size of the genome-wide association studies (GWAS) used to compute the PGS

a) Lee et al. et al. (2018, N = 1,131,881 (1)

b) Okbay et al. et al. (2016, N = 293,723 (0.5)

c) Rietveld et al. et al. (2013, N = 101,069 (0)

4) Sample overlap

a) the cohort does not overlap with the sample used to derive the GWAS (1)

b) an updated GWAS excluding the cohort is used (1)

c) the cohort is used to derive the GWAS (0)

5) Genetic ancestry

a) the cohort is from the same genetic ancestry with the GWAS[2] (1)

b) the cohort is from different genetic ancestry with the GWAS[2] (0)

[2]European descent

Comparability/confounding

6) Study fully accounts for genetic nurture effects

a) yes, study uses control by design, i.e., virtual parent design (1)

b) yes, study uses statistical control and accounted for PGS of both parents and child (1)

c) yes, study uses statistical control and accounted for PGS of one parent and child (0.5)

d) no (0)

7) Study accounts for the confounding of age, sex and principal components (PC)

a) yes, study controls for both age, sex and PC (1)

b) yes, study controls for sex and PC and all participants were the same on age (1)

c) yes, study partly controls for age, sex and PC (0.5)

d) no (0)

Outcome: Educational attainment and educational achievement

8) Assessment of outcome

a) official record (1)

b) instrument tested for validity and reliability (1)

c) self-report (0.5)

b) no description (0)

9) Same underlying phenotype of outcome and GWAS

a) the outcome represents the same underlying phenotype as measured by the GWAS used, e.g., years of education, highest degree obtained (1)

b) the outcome represents somewhat the same underlying phenotype as measured by the GWAS used, e.g., national tests scores, school grades (0.5)

c) the outcome presents different phenotype as measured by the GWAS used, e.g., cognitive performance, math course level, EA difference between twins (0)

Detailed methodological score of each included study see sTable 4. It should be noted that most of the included studies (9 out of 12) had limited representativeness of their original cohort. Taking the minimal attrition rate per study on genetic nurture estimates, the median attrition rate was 55.6% (attrition rate for each estimate see sTable 3). This substantial rate in attrition among the included studies is mainly due to missingness of genetic data (i.e. samples not genotyped within cohorts) or ancestry restriction due to predominantly European-descent samples of GWAS (McGuire et al., 2020).

### 4. Data extraction and synthesis

#### 4.1 Effect size calculation

Standardized beta coefficients, which constituted the most commonly reported metric among the included studies, were used to measure effect sizes. One study (Conley et al., 2015) reported unstandardized betas, which were manually standardized by multiplying unstandardized coefficients by the ratio of standard deviations of the corresponding independent variable and dependent variable. Some studies did not report the standard error of estimates, which was necessary for estimating the pooled effect size. For studies that reported 95% confidence intervals (Armstrong-Carter et al., 2020; Bates et al., 2019; Bates et al., 2018; de Zeeuw et al., 2020; Wertz et al., 2019), standard errors were manually calculated by dividing corresponding confidence intervals (upper limit – lower limit) by 3.92. For studies that reported t-test statistics (Belsky et al., 2016), standard errors were manually calculated by dividing corresponding standardized betas by t-test statistics. For studies that did not report data allowing for direct calculation of standard errors (Kong et al., 2018), following non-response after contacting the author, we imputed standard errors from corresponding estimates and sample sizes by using the compute.es_0.2-4 package (Del Re, 2013) in R version 3.6.1 (R Core Team, 2019).

Genetic nurture effects in one of the included studies (Willoughby, McGue, Iacono, Rustichini, & Lee, 2019) using average parental PGS was recalibrated to improve comparability with other studies using individual parental PGS. Details see in section 7.2.

For studies using the virtual parent design (Bates et al., 2019; Bates et al., 2018; de Zeeuw et al., 2020; Kong et al., 2018), estimates of unadjusted child effects (based on transmitted PGS) and genetic nurture effects (based on non-transmitted PGS) were reported. The magnitude of direct genetic effects can be imputed as follows:

In regressions of the child education on both parental transmitted polygenic score ($PGS_T$) and non-transmitted polygenic score ($PGS_{NT}$), let $N$ be the sample size, $\beta_T$ be the coefficient of $PGS_T$ and $\beta_{NT}$ the coefficient of $PGS_{NT}$, $\sigma_T$ be the standard error of $\beta_T$ and $\sigma_{NT}$ be the standard error of $\beta_{NT}$. $\beta_T$ is the effect of the children's PGS on their own phenotype and $\beta_{NT}$ is the effect of genetic nurture. Thus, estimated direct genetic effects $\beta_{direct} = \beta_T - \beta_{NT}$ (Kong et al., 2018). The variance of $\beta_{direct}$ is equal to the sum of variance of $\beta_T$ and $\beta_{NT}$. Let $\sigma_{direct}$ denotes the standard error of direct genetic effects, then it can be expressed as: $\sigma_{direct} = \sqrt{(\sigma_T^2 + \sigma_{NT}^2)}$.

#### 4.2 Multilevel Random Effects Model (MREM)

The three-level MREM was performed following the tutorial of Assink and Wibbelink (Assink & Wibbelink, 2016) and incorporated variation in effect sizes from three sources: Level 1 variance attributed to random sampling, which corresponds to the standard error of each individual effect size; Level 2 variation between estimates within a single cohort; Level 3 variation in estimates between different cohorts. Here the cohort level was defined as the original population-based sample on which the data about genotype and educational outcomes was collected. In this meta-analysis, we included 12 studies from eight independent cohorts (see Table 1, column 'Cohort'). This level was employed to account for the violation of independence due to multiple estimates and studies from the same cohort. For example, two studies (Conley et al., 2015; Liu, 2018) were both from the Framingham Heart Study (FHS), and in one studies (Liu, 2018) several estimates of genetic nurture effects were reported using genotype from mothers and fathers. One special case was that participants in the Minnesota Center for Twin and Family Research (MCTFR) cohort were drawn from several longitudinal studies including the Minnesota Twin Family Study (MTFS) cohort, thus in the meta-analysis they were considered as the same cohort.

##### 4.3 Moderator analyses of study characteristics

We tested moderating roles of a number of study characteristics reflecting methodological quality and sample representativeness, including study quality, sample size and attrition rate of the cohort. Study quality was indexed by the total score of the methodological quality described in section 3.2. Detailed score of each included study see sTable 4. The sample size was used in the unit of 1,000 participants due to the relatively large sample size in studies (mean = 3,372, median = 1,626). For the sample size of each effect size, see sTable 3. To note, only the largest sample size assessing genetic nurture effects in each cohort was used to compute the total sample size of the current meta-analysis (i.e., 38,654) in a conservative manner to preclude any overlap within the cohort. In Table1, sample sizes were reported per study and outcome category (i.e., educational attainment vs. educational achievement) as part of the study summary. Considering the prevalent attrition in original cohorts across studies (details see Section 3.3), the moderating role of attrition rate was tested. For the attrition rate of each effect size, see sTable 3.

#### 5. Unadjusted parental and child effects

##### 5.1 Unadjusted parental effects

We derived k = 8 estimates of unadjusted parental effects on offspring educational outcomes (i.e., without adjusting for genetic nurture effects). Table 2 shows MREM findings when jointly meta-analysing genetic nurture effects and unadjusted parental effects ($\beta_{\text{mixture}}$ = 0.11, 95% CI [0.08, 0.14], robust CI [0.07, 0.14]). When evaluating the heterogeneity in effect sizes, we found that a substantial proportion of variance reflected within-cohort heterogeneity ($I^2_{\text{Level 2}}$ = 65.83%). This indicates that within-cohort factors (i.e., differences in effect sizes between outcomes within the same cohort) may account for some of the variation in effect sizes. Results of subgroup analysis showed that the magnitude of genetic nurture effects and unadjusted parental effects were highly significantly different ($Q = 20.58$, $df = 1$, $p < .0001$). As is shown in Table 2 and Figure 2, estimates from unadjusted parental effects ($\beta_{\text{parental unadjusted}} = 0.21$, 95% CI [0.15, 0.26], robust CI [0.08-0.33]) only were larger than genetic nurture. The variance among effect sizes of unadjusted parental effects mainly resulted from the between-cohort heterogeneity ($I^2_{\text{Level 3}} = 86.86\%$), suggesting that factors in which the cohorts may differ (e.g., type of the educational outcome, age when the outcome was assessed, accuracy of the GWAS) accounted for some of the variation in effect sizes. The funnel plot (see sFigure 1) and formal test with precision as a moderator ($Q = 3.22$, $p = .0727$) suggested no publication bias in estimates of unadjusted parental effects. Results of jack-knife leave-one-out analysis (see sFigure 2) suggest no substantial role of a single influential study. Results of moderation analyses for unadjusted parental PGS are shown in sTable 6. Studies using PGS derived from more accurate GWAS reported larger unadjusted parental effects ($p_{\text{unadjusted parent}} < .0001$). Moreover, the type of educational outcome ($p < .0001$) and study quality score ($p = .0353$) explained some variation in the effect sizes.

##### 5.2 Unadjusted child effects

We derived k = 11 estimates of unadjusted child effects on their own educational outcomes, i.e., effects of child PGS without considering genetic nurture effects. Table 2 shows MREM findings of child effects on education when jointly meta-analysing direct genetic effects and unadjusted child effects ($\beta_{\text{mixture}} = 0.20$, 95% CI [0.16, 0.24], robust CI [0.15, 0.24]). More variance among effect sizes in the joint model was attributable to between-cohort heterogeneity than within-cohort heterogeneity ($I^2_{\text{Level 2}} = 36.17\%$ versus $I^2_{\text{Level 3}} = 55.13\%$). Results of subgroup analysis showed that magnitudes of direct genetic effects and unadjusted child effects were significantly different ($Q = 6.39$, $df = 1$, $p = .0115$). As is shown in Table 2 and Figure 3, the magnitude of unadjusted child effects ($\beta_{\text{child unadjusted}}$ = 0.24, 95% CI [0.20, 0.28], robust CI [0.19, 0.29]) was larger than direct genetic effects. More variance among effect sizes in unadjusted child effects was attributable to between-cohort heterogeneity than within-cohort heterogeneity ($I^2_{\text{Level 2}}$ = 28.51% versus $I^2_{\text{Level 3}}$ = 59.92%).The funnel plot (see sFigure 3) and formal test with precision as a moderator suggested no publication bias in estimates of unadjusted child effects ($Q$ = 0.47, $p$ = 0.4917). Results of jack-knife leave-one-out analysis (see sFigure 4) suggests no substantial role of a single influential study. Unadjusted child effects were consistent across most of tested moderators, except that studies using PGS derived from more accurate GWAS reported larger effect sizes ($p_{\text{unadjusted child}}$ = .0010).

#### 6. Family-level adjustment

We tested the moderating role of family-level adjustment, including parental education level and family socioeconomic status (SES) to quantify the extent to which genetic nurture effects can be attributed to these distal family-level factors. We compared effect sizes with and without family-level adjustments. Effect sizes included in the main meta-analysis were unadjusted for family-level adjustment ($k_{\text{genetic nurture}}$ =22, $k_{\text{direct genetic}}$ = 16, $k_{\text{parental unadjusted}}$ =8, $k_{\text{child unadjusted}}$ =11). All available effect sizes in the included studies with family-level adjustment of parental education or family SES were extracted as adjusted estimates ($k_{\text{genetic nurture}}$ =18, $k_{\text{direct genetic}}$ = 11, $k_{\text{parental unadjusted}}$ =4, $k_{\text{child unadjusted}}$ =3). Effect sizes adjusted for family-level factors were only used to test the moderating role of family-level adjustment and not included in the main meta-analysis as they were fundamentally different from unadjusted ones. It should be noted that one study (Conley et al., 2015) reported genetic nurture and direct genetic effects only with family-level adjustment, and unadjusted parental and child effects both with and without family-level adjustment. Therefore, for this particular study

(Conley et al., 2015), only effect sizes of unadjusted parental and child effects were included in the main meta-analysis, and all effect sizes were used for the moderator analysis of family-level adjustment.

The unadjusted effects were visually larger than the family-level adjusted effects for both genetic nurture and direct genetic effects. The largest decrease in effect sizes attributable to adjustment was present for genetic nurture effects ($\beta_{\text{unadjusted}}$ = 0.07, 95% CI [0.07, 0.08] vs. $\beta_{\text{adjusted}}$ = 0.02, 95% CI [0.01, 0.03]), which was supported by a highly significant moderating effect ($p_{\text{adjustment}} < .0001$) and remained robust when tested in sensitivity analysis (see sTable 7). Smaller changes in effect sizes following family-level adjustment were present for direct genetic effects ($\beta_{\text{unadjusted}}$ = 0.17, 95% CI [0.13, 0.20] vs. $\beta_{\text{adjusted}}$ = 0.14, 95% CI [0.10, 0.18]), in which case the moderating effect was also significant ($p_{\text{adjustment}} = .0098$).

After accounting for parental education level or family SES, the effect of unadjusted parental and child effects on children's educational outcomes were both attenuated by ~30%. ($p_{\text{unadjusted parent}} = .0001$, $p_{\text{unadjusted child}} = .0223$).

### 7. Sensitivity analyses

#### 7.1 Robust confidence intervals of dependent estimates

Among some included studies, we extracted multiple, statistically dependent effect size estimates from the same cohort. We utilize MREM to handle the dependence of effect sizes. As sensitivity checks, we also reported, robust confidence intervals (robust CI) of cluster-robust variance estimations, which is out of the MREM framework and obtained using the package clubSandwich version 0.5.0 (Pustejovsky, 2020) in R version 3.6.1 (R Core Team, 2019).

#### 7.2 Impact of recalibrating estimates using the average parental PGS

In general, comparing estimates using maternal, paternal, maternal and/or paternal genomic measures should be straightforward by directly compared their absolute values. Caution is warranted when using the average parental PGS, in which case the $R^2$ (variance explained) is unbiased but the estimate is inflated (compared to using individual parental PGS):

Let $PGS_M$ be the maternal polygenic score and $PGS_P$ be the paternal polygenic score, and let $R^2_M$ and $R^2_P$ be the variance explained by $PGS_M$ and $PGS_P$, respectively. The variance explained by the average parental PGS, $R^2_{ave\ parent}$, equals to the addition of variance of mother and father ($R^2_{ave\ parent} = R^2_M + R^2_P$) when assuming $PGS_M$ and $PGS_P$ are uncorrelated. Thus, the standardized estimated genetic nurture effects from the average parental PGS, $\beta_{ave\ parent}$ is equal to the square root of $R^2_M + R^2_P$. Assuming that genetic nurture effects from mother and father are equal ($R^2_M = R^2_P = R^2_{ind\ parent}$), and let $\beta_{ind\ parent}$ be the standardized estimate of genetic nurture effects from the individual parental PGS, the relationship between $\beta_{ave\ parent}$ and $\beta_{ind\ parent}$ are:

Genetic nurture effects from the average parental PGS:

$$\beta_{ave\ parent} = \sqrt{(2R^2_{ind\ parent})} = \sqrt{2}\,\beta_{ind\ parent}$$

Genetic nurture effects from the individual parental PGS:

$$\beta_{ind\ parent} = \sqrt{(R^2_{ave\ parent}/2)} = \beta_{ave\ parent}/\sqrt{2}$$

Due to the abovementioned reason, one of the included studies (Willoughby et al., 2019) utilizing the average parental PGS to capture genetic nurture effects had an outlying estimate ($\beta_{Willoughby\ original} = 0.20$) relative to other estimates included in our study. We thus used the recalibrated estimate ($\beta_{Willoughby\ adjusted} = 0.20/\sqrt{2} = 0.14$) for the main meta-analysis to obtain better comparability with other studies using individual parental PGS.

The impact of this adjustment is examined by meta-analysing genetic nurture effects with the originally reported effect size. Using the original estimate of average parental PGS resulted in similar genetic nurture effects ($\beta_{\text{Willoughby original}}$ = 0.08, 95% CI [0.07, 0.09], robust CI [0.06, 0.10]) but introduced more publication bias ($Q$ = 8.59, $p$ =.0034). The moderating effect of analytical design was still significant ($Q$ = 6.17, $p$ = 0.0457), with smaller estimates from the virtual parent design than the statistical control approach. Such results were expected, as the adjusted average parental PGS was derived from the statistical control approach and the recalibration decreased that estimate.

To note, when our pooled estimate of genetic nurture effects from the individual parental PGS is reversely adjusted to represent genetic nurture effects from the average PGS of both parents, $\beta_{\text{ave parent}} = \sqrt{2}\ \beta_{\text{ind parent}} = \sqrt{2}*0.08 = 0.11$. The ratio of genetic nurture effects and direct genetic effects increases to 0.11/0.17 = 0.65. This finding is comparable to a recent study using Relatedness Disequilibrium Regression (Young et al., 2018). In Young et al. 2018, genetic nurture effects explained 6.6% of the variance in educational attainment (the effect size is thus $\sqrt{0.066}$ = 0.26) and direct genetic effects/heritability explained 17% of the variance in educational attainment (the effect size is thus $\sqrt{0.17}$ = 0.41), resulting in a ratio 0.63 (i.e., 0.26/0.41).

##### 7.3 Impact of a potentially influential study

Due to the Inverse-variance weighting strategy of meta-analysis, one of the included studies (Kong et al., 2018) might be more influential than others since its standard errors (imputed based on their corresponding effect sizes and sample sizes) were very small. Therefore, we tested the impact of this potentially influential study by re-running the meta-analysis omitting its estimates. We tested the robustness of our pooled effects as well as the distribution of variance in the MREM to see whether the narrow confidence intervals and approximate homogeneity of genetic nurture effects were exclusively attributed to this study, e.g., the Kong et al. 2018 study reported three estimates of genetic nurture effects using maternal, paternal and parental non-transmitted PGS. In addition, we performed meta-regression without estimates from this study for all the moderators that were potentially impacted, since the Kong et al. 2018 study may have independently influenced the moderating effects in some cases. For example, multiple genetic nurture effects from the Kong et al. 2018 study using maternal, paternal and parental non-transmitted PGS) may unduly impact on the moderating effect of parent of origin and mask effects from other studies.

The potentially influential study of Kong et al. 2018 did not show substantial impact on distributions of variance, pooled estimates, but resulted in some publication bias and changes in moderating effects of methodological quality and sample size on the magnitude of genetic nurture effects. For details see sTables 5 and 7.

##### 7.4 Jack-knife leave-one-out analyses

To account for any other potential influences from included studies, we assessed the undue effect of individual studies on our pooled estimates through jack-knife leave-one-out analyses, by testing changes in the estimate across permutations in which each study was omitted in turn. For visualization of results see sFigures 2 and 4.

##### 7.5 The moderating effect of outcome type within study

Two of the included studies (de Zeeuw et al., 2020; Rustichini, Iacono, Lee, & McGue, 2018) assessed both educational attainment and achievement, we thus checked the robustness of moderating effect of outcome type within study by running the meta regression within these two particular studies. The moderation role for outcome type on genetic nurture effects became statistically significant ($\beta_{\text{attainment}}$ = 0.11, 95% CI [0.08, 0.14], $\beta_{\text{achievement}}$ = 0.05, 95% CI [0.01, 0.09], $p$ = .0228). The difference in direct genetic effects was larger in the opposite direction ($\beta_{attainment}$ = 0.12, 95% CI [0.07, 0.17]), $\beta_{achievement}$ = 0.21, 95% CI [0.16, 0.26], $p$ = .0144).

Several plausible explanations might account for the consistently higher heritability/direct genetic effect in educational achievement. One is that educational achievement is measured during compulsory schooling. The difference thus may reflect more genetically influenced traits in children like intelligence, personality and psychopathology (Krapohl et al., 2014). In contrast, years of education one completed is likely to be influenced by a wider range of factors, such as career plan or financial situation (Abdellaoui et al., 2019; Harden et al., 2020; Morris, Davies, & Smith, 2020). Another explanation is that educational achievement is often measured with standard tests/scores which reflect one's relative ranking/decile among peers. Education years, however, can be more ambiguous as essentially different routes, such as academic and vocational, are not distinguished, which may introduce heterogeneity and measurement error. Future studies should examine genomic predictions with consistent measures of educational achievement across different developmental stages in order to capture dynamic changes. Evidence from one study, the Twins Early Development Study (TEDS), suggested that the predictive precision of PGS (i.e., unadjusted child effects) on educational achievement increased from ages 7 to 16 (Selzam et al., 2017).

### 8. sFigures

#### 8.1 sFigure 1. Funnel plots for parental effects on educational outcomes

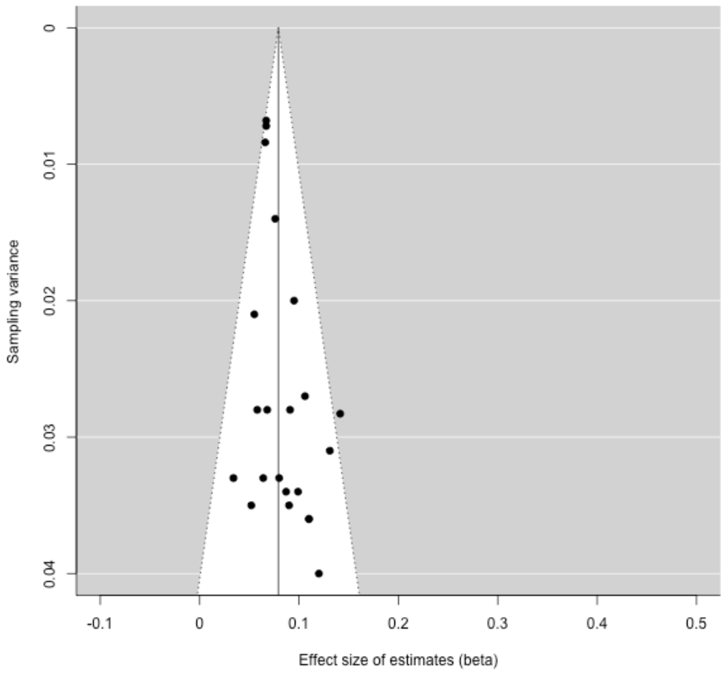


A. Genetic nurture effects

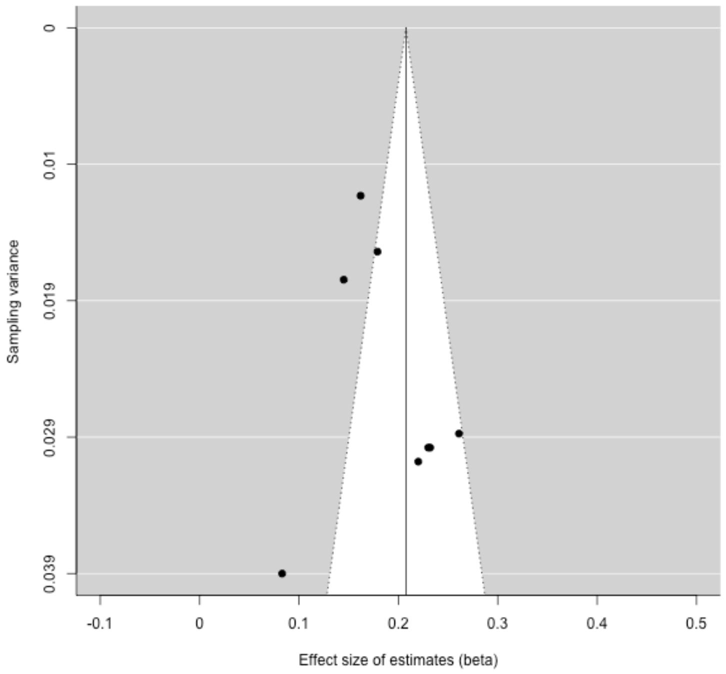


B. Unadjusted parental effects.

#### 8.2 sFigure 2. Jackknife sensitivity analyses for parental effects on educational outcomes

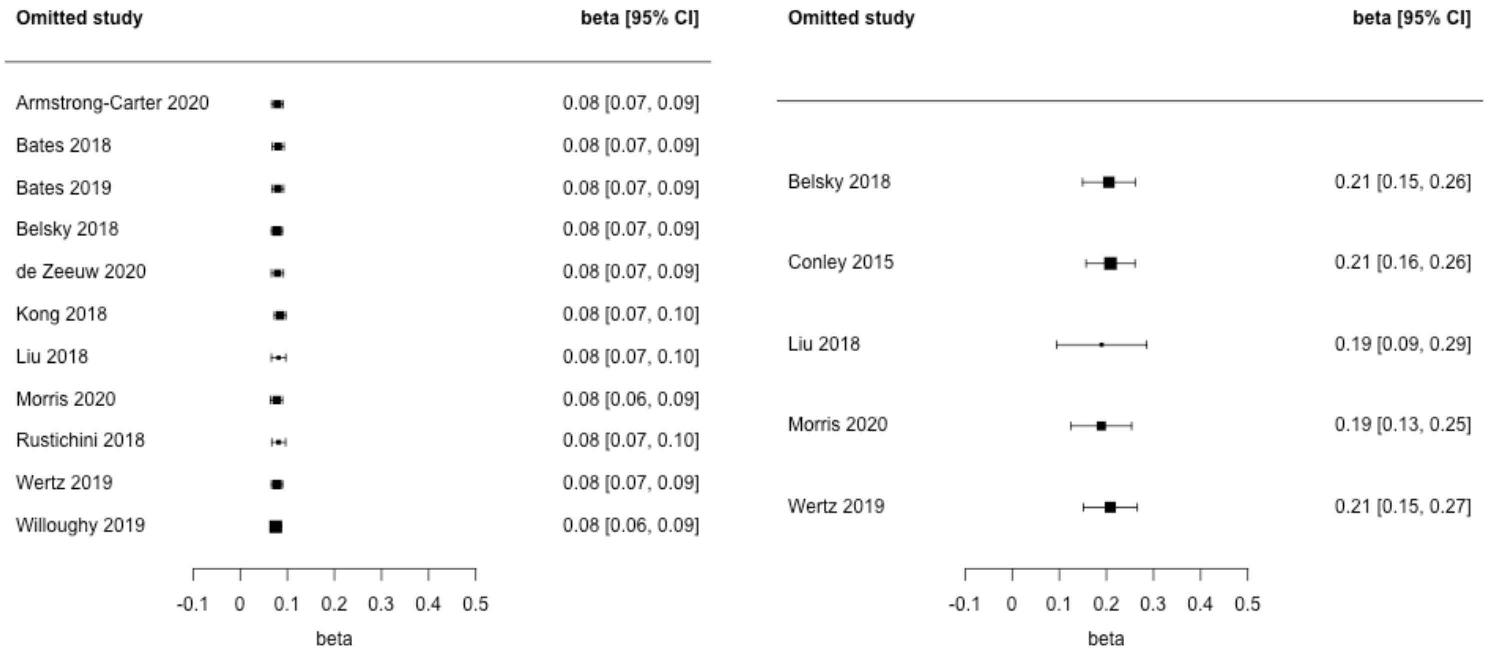


A. Genetic nurture effects

B. Unadjusted parental effects.

The estimate corresponding to each study listed reflects the pooled beta from a meta-analysis where that study was omitted.

#### 8.3 sFigure 3. Funnel plots for child effects on educational outcomes

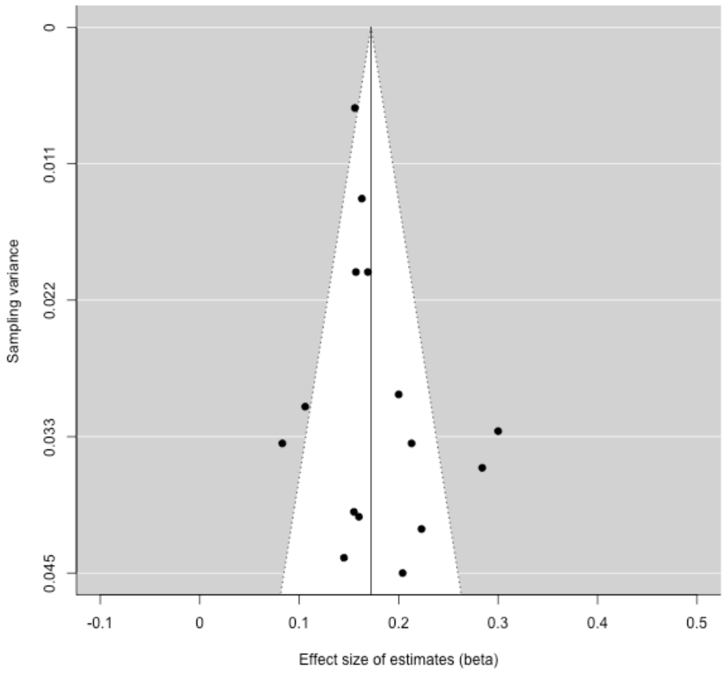


A. Direct genetic effects

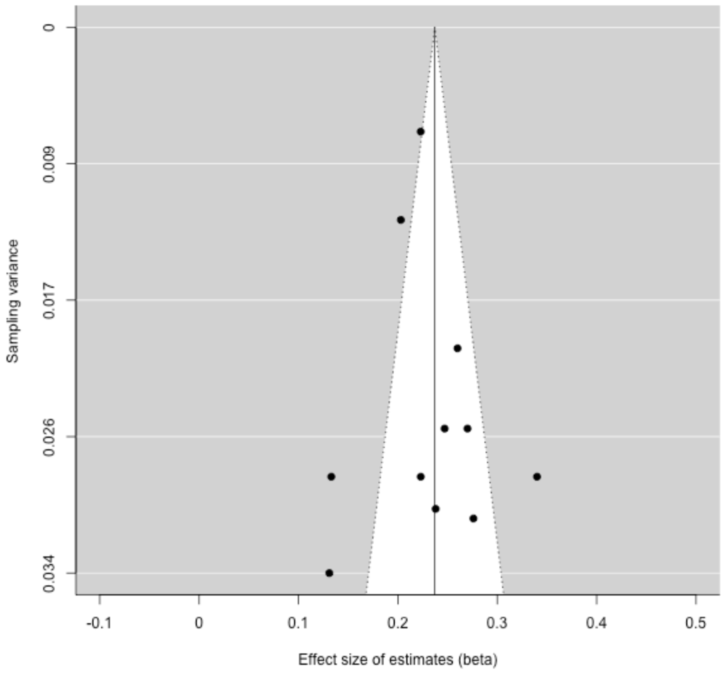


B. Unadjusted child effects.

#### 8.4 sFigure 4. Jackknife sensitivity analyses for child effects on educational outcomes

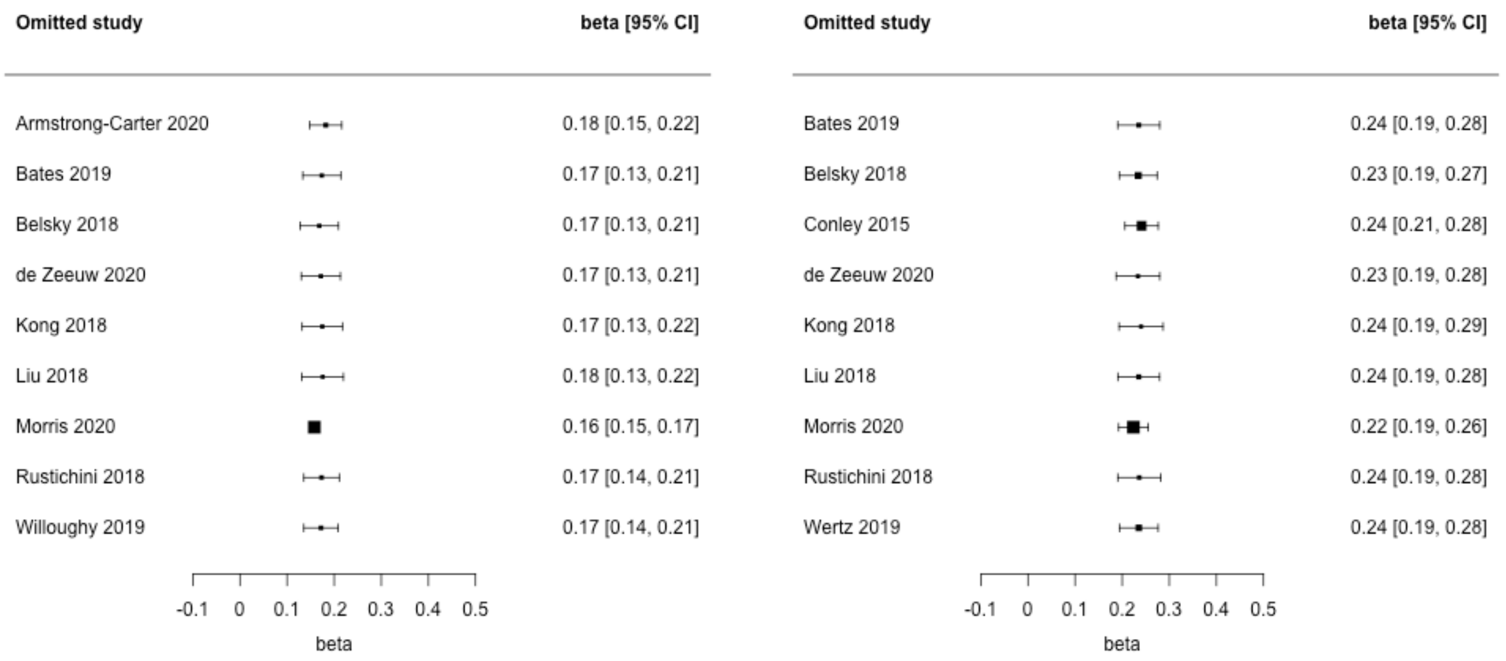


A. Direct genetic effects B. Unadjusted child effects

The estimate corresponding to each study listed reflects the pooled beta from a meta-analysis where that study was omitted.

### 9. sReferences


Abdellaoui, A., Hugh-Jones, D., Yengo, L., Kemper, K. E., Nivard, M. G., Veul, L., . . . Wray, N. R. (2019). Genetic correlates of social stratification in Great Britain. *Nature human behaviour, 3*(12), 1332-1342.

Armstrong-Carter, E., Trejo, S., Hill, L. J. B., Crossley, K. L., Mason, D., & Domingue, B. W. (2020). The Earliest Origins of Genetic Nurture: The Prenatal Environment Mediates the Association Between Maternal Genetics and Child Development. *Psychological science*. Retrieved from <Go to ISI>://WOS:000537515200001

Assink, M., & Wibbelink, C. J. (2016). Fitting three-level meta-analytic models in R: A step-by-step tutorial. *The Quantitative Methods for Psychology, 12*(3), 154-174.

Bates, T. C., Maher, B. S., Colodro-Conde, L., Medland, S. E., McAloney, K., Wright, M. J., . . . Gillespie, N. A. (2019). Social competence in parents increases children's educational attainment: Replicable genetically-mediated effects of parenting revealed by non-transmitted DNA. *Twin Research and Human Genetics, 22*(1), 70-74. Retrieved from http://journals.cambridge.org/action/displayJournal?jid=THG http://ovidsp.ovid.com/ovidweb.cgi?T=JS&PAGE=reference&D=emexa&NEWS=N&AN=626076119

Bates, T. C., Maher, B. S., Medland, S. E., McAloney, K., Wright, M. J., Hansell, N. K., . . . Gillespie, N. A. (2018). The nature of nurture: Using a virtual-parent design to test parenting effects on children's educational attainment in genotyped families. *Twin Research and Human Genetics, 21*(2), 73-83.

Belsky, D. W., Moffitt, T. E., Corcoran, D. L., Domingue, B., Harrington, H., Hogan, S., . . . Williams, B. S. (2016). The genetics of success: How single-nucleotide polymorphisms associated with educational attainment relate to life-course development. *Psychological science, 27*(7), 957-972.

Chabris, C. F., Lee, J. J., Cesarini, D., Benjamin, D. J., & Laibson, D. I. (2015). The fourth law of behavior genetics. *Current Directions in Psychological Science, 24*(4), 304-312.

Cheesman, R., Hunjan, A., Coleman, J. R., Ahmadzadeh, Y., Plomin, R., McAdams, T. A., . . . Breen, G. (2020). Comparison of adopted and nonadopted individuals reveals gene–environment interplay for education in the UK Biobank. *Psychological science, 31*(5), 582-591.

Conley, D., Domingue, B. W., Cesarini, D., Dawes, C., Rietveld, C. A., & Boardman, J. D. (2015). Is the Effect of Parental Education on Offspring Biased or Moderated by Genotype? *Sociological science, 2*(6), 82-105. Retrieved from http://ovidsp.ovid.com/ovidweb.cgi?T=JS&PAGE=reference&D=prem1&NEWS=N&AN=29051911

de Zeeuw, E. L., Hottenga, J. J., Ouwens, K. G., Dolan, C. V., Ehli, E. A., Davies, G. E., . . . van Bergen, E. (2020). Intergenerational Transmission of Education and ADHD: Effects of Parental Genotypes. *Behavior Genetics*. doi:10.1007/s10519-020-09992-w

Del Re, A. C. (2013). compute.es: Compute Effect Sizes. Retrieved from https://cran.r-project.org/package=compute.es

Domingue, B. W., & Fletcher, J. (2020). Separating measured genetic and environmental effects: Evidence linking parental genotype and adopted child outcomes. *Behavior Genetics*, 1-9.

Dudbridge, F. (2013). Power and predictive accuracy of polygenic risk scores. *PLoS Genet, 9*(3), e1003348.

Gratten, J., Wray, N. R., Keller, M. C., & Visscher, P. M. (2014). Large-scale genomics unveils the genetic architecture of psychiatric disorders. *Nature Neuroscience, 17*(6), 782-790. doi:10.1038/nn.3708

Harden, K. P., Domingue, B. W., Belsky, D. W., Boardman, J. D., Crosnoe, R., Malanchini, M., . . . Harris, K. M. (2020). Genetic associations with mathematics tracking and persistence in secondary school. *NPJ science of learning, 5*(1), 1-8.

International Schizophrenia Consortium. (2009). Common polygenic variation contributes to risk of schizophrenia that overlaps with bipolar disorder. *Nature, 460*(7256), 748.

Kong, A., Thorleifsson, G., Frigge, M. L., Vilhjalmsson, B. J., Young, A. I., Thorgeirsson, T. E., . . . Masson, G. (2018). The nature of nurture: Effects of parental genotypes. *Science, 359*(6374), 424-428.

Krapohl, E., Rimfeld, K., Shakeshaft, N. G., Trzaskowski, M., McMillan, A., Pingault, J.-B., . . . Plomin, R. (2014). The high heritability of educational achievement reflects many genetically influenced traits, not just intelligence. *Proceedings of the national academy of sciences, 111*(42), 15273. doi:10.1073/pnas.1408777111

Lee, J. J., Wedow, R., Okbay, A., Kong, E., Maghzian, O., Zacher, M., . . . Linnér, R. K. (2018). Gene discovery and polygenic prediction from a 1.1-million-person GWAS of educational attainment. *Nature Genetics, 50*(8), 1112.

Liu, H. X. (2018). Social and Genetic Pathways in Multigenerational Transmission of Educational Attainment. *American sociological review, 83*(2), 278-304. Retrieved from <Go to ISI>://WOS:000429752000003

McGuire, A. L., Gabriel, S., Tishkoff, S. A., Wonkam, A., Chakravarti, A., Furlong, E. E. M., . . . Kim, J.-S. (2020). The road ahead in genetics and genomics. *Nature Reviews Genetics, 21*(10), 581-596. doi:10.1038/s41576-020-0272-6

Morris, T. T., Davies, N. M., & Smith, G. D. (2020). Can education be personalised using pupils' genetic data? *eLife, 9*, e49962.

Okbay, A., Beauchamp, J. P., Fontana, M. A., Lee, J. J., Pers, T. H., Rietveld, C. A., . . . Meddens, S. F. W. (2016). Genome-wide association study identifies 74 loci associated with educational attainment. *Nature, 533*(7604), 539-542.

Ouzzani, M., Hammady, H., Fedorowicz, Z., & Elmagarmid, A. (2016). Rayyan—a web and mobile app for systematic reviews. *Systematic Reviews, 5*(1), 210. doi:10.1186/s13643-016-0384-4

Pustejovsky, J. (2020). clubSandwich: Cluster-Robust (Sandwich) Variance Estimators with Small-Sample Corrections.

R Core Team. (2019). R: A language and environment for statistical computing. Vienna, Austria: R Foundation for Statistical Computing.

Rietveld, C. A., Medland, S. E., Derringer, J., Yang, J., Esko, T., Martin, N. W., . . . Agrawal, A. (2013). GWAS of 126,559 individuals identifies genetic variants associated with educational attainment. *Science, 340*(6139), 1467-1471.

Ronald, A. (2020). Polygenic scores in child and adolescent psychiatry–strengths, weaknesses, opportunities and threats. *Journal of Child Psychology and Psychiatry, 61*(5), 519-521.

Rustichini, A., Iacono, W. G., Lee, J., & McGue, M. (2018). *Polygenic score analysis of educational achievement and intergenerational mobility*.

Selzam, S., Krapohl, E., von Stumm, S., O'Reilly, P., Rimfeld, K., Kovas, Y., . . . Plomin, R. (2017). Predicting educational achievement from DNA. *Molecular Psychiatry, 22*(2), 267-272.

Selzam, S., Ritchie, S. J., Pingault, J.-B., Reynolds, C. A., O'Reilly, P. F., & Plomin, R. (2019). Comparing within-and between-family polygenic score prediction. *The American Journal of Human Genetics, 105*(2), 351-363.

Shen, H., & Feldman, M. W. (2020). Genetic nurturing, missing heritability, and causal analysis in genetic statistics. *Proceedings of the national academy of sciences, 117*(41), 25646-25654.

Stang, A. (2010). Critical evaluation of the Newcastle-Ottawa scale for the assessment of the quality of nonrandomized studies in meta-analyses. *European journal of epidemiology, 25*(9), 603-605.

Wertz, J., Moffitt, T. E., Agnew-Blais, J., Arseneault, L., Belsky, D. W., Corcoran, D. L., . . . Caspi, A. (2019). Using DNA From Mothers and Children to Study Parental Investment in Children's Educational Attainment. *Child development*. Retrieved from http://ovidsp.ovid.com/ovidweb.cgi?T=JS&PAGE=reference&D=emexa&NEWS=N&AN=629707797

Willoughby, E. A., McGue, M., Iacono, W. G., Rustichini, A., & Lee, J. J. (2019). The role of parental genotype in predicting offspring years of education: evidence for genetic nurture. *Molecular Psychiatry*. Retrieved from http://www.nature.com/mp/index.html http://ovidsp.ovid.com/ovidweb.cgi?T=JS&PAGE=reference&D=emexa&NEWS=N&AN=2002635365

Wray, N. R., Goddard, M. E., & Visscher, P. M. (2007). Prediction of individual genetic risk to disease from genome-wide association studies. *Genome research, 17*(10), 1520-1528.

Young, A. I., Frigge, M. L., Gudbjartsson, D. F., Thorleifsson, G., Bjornsdottir, G., Sulem, P., . . . Kong, A. (2018). Relatedness disequilibrium regression estimates heritability without environmental bias. *Nature Genetics, 50*(9), 1304-1310. doi:10.1038/s41588-018-0178-9
